# *TTBK2* mutations associated with spinocerebellar ataxia type 11 disrupt peroxisome dynamics and ciliary localization of SHH signaling proteins

**DOI:** 10.1101/2023.01.31.526333

**Authors:** Jesús Muñoz-Estrada, Abraham V. Nguyen, Sarah C. Goetz

## Abstract

Frameshift mutations in *Tau Tubulin Kinase 2* (*TTBK2*) cause spinocerebellar ataxia type 11 (SCA11), which is characterized by the progressive loss of Purkinje cells and cerebellar atrophy. Previous work showed that these *TTBK2* variants generate truncated proteins that interfere with primary ciliary trafficking and with Sonic Hedgehog (SHH) signaling in mice. Nevertheless, the molecular mechanisms underlying the dominant interference of mutations remain unknown. Herein, we discover that SCA11-associated variants contain a *bona fide* peroxisomal targeting signal type 1. We find that their expression in RPE1 cells reduces peroxisome numbers within the cell and at the base of the cilia, disrupts peroxisome fission pathways, and impairs trafficking of ciliary SMO upon SHH signaling activation. This work uncovers a neomorphic function of SCA11-causing mutations and identifies requirements for both peroxisomes and cholesterol in trafficking of cilia-localized SHH signaling proteins. In addition, we postulate that molecular mechanisms underlying cellular dysfunction in SCA11 converge on the SHH signaling pathway.

**SUMMARY:** Molecular mechanisms underlying spinocerebellar ataxia type 11 are not well understood. In this study, we identified a neomorphic function of the mutated gene (*TTBK2*) associated with this disease highlighting a functional inter-organelle interaction between peroxisomes and cilia.

## INTRODUCTION

*TTBK2* gene encodes for Tau Tubulin Kinase 2 (TTBK2) protein, a serine/threonine kinase that localizes at the distal end of the mother centriole at structures known as the distal appendages (DAs) (Goetz et al., 2012; Tanos et al., 2013). In quiescent and non-proliferating cells, including neurons, its catalytic activity at the DAs is required for the assembly of a functional primary cilium (Bernatik et al., 2020; Cajanek and Nigg, 2014; Goetz et al., 2012). The lack of this kinase leads to disruption of Sonic hedgehog (SHH) cilia-dependent processes during embryonic development and the connectivity and integrity of Purkinje cells in the adult cerebellum (Bowie and Goetz, 2020; Bowie et al., 2018).

Primary cilia are non-motile microtubule-based structures that project from the cell surface and are enriched in receptors important for SHH and GPCR signaling in a wide variety of cell types, including differentiated neurons (Bansal et al., 2019; Hsiao et al., 2021; Stubbs et al., 2022). Cilia are surrounded by a membrane with a unique protein and lipidic composition and its compartmentalization from the cytoplasm is regulated by a domain proximal to its base known as ciliary transition zone (Garcia-Gonzalo and Reiter, 2017). Despite our knowledge of the molecular mechanisms maintaining ciliary receptor localization and trafficking (Mukhopadhyay et al., 2017; Nachury and Mick, 2019), less progress has been made towards understanding how the lipidic composition of these structures is regulated. A recent study reported that cholesterol at the ciliary membrane is trafficked by a peroxisome-associated protein complex (Miyamoto et al., 2020). Furthermore, a current model for SHH signaling regulation indicate the importance of a specific pool of cholesterol in the outer leaflet of the plasma membrane (Kinnebrew et al., 2021).

Mutations in genes that regulate cilia formation and function cause clinically heterogenous human developmental disorders collectively termed ciliopathies. These disorders are characterized by malformation and dysfunction of multiple organ systems, including neurological features such as cerebellar malformations and various levels of intellectual disability (Reiter and Leroux, 2017). On the other hand, mutations in TTBK2 are reported to cause autosomal dominant spinocerebellar ataxia type11 (or SCA11) (Houlden et al., 2007). SCA11 is a rare movement disorder characterized by loss of Purkinje cells in the cerebellum. Currently, five different mutations in *TTBK2* have been reported in affected families with SCA11 worldwide. Four are frameshift mutations that originate a premature stop codon and truncated proteins of approximately 450 amino acids (Bauer et al., 2010; Houlden et al., 2007; Lindquist et al., 2017) and a recently reported missense mutation in its C-terminal domain (p.Val1097Ala) (Deng et al., 2020). Previous work by our lab showed dominant interference in cilia formation and SHH signaling by a frameshift SCA11 mutation (or *TTBK2*^fam1^ mutation) in mice (Bowie et al., 2018). Intriguingly, TTBK2^fam1^ protein do not interact with full-length TTBK2 protein and therefore, the molecular mechanism of how SCA11-associated variants interfere with cilia function remains unsolved (Bowie et al., 2018). To address this question, we performed biotin proximity labeling with both TTBK2^WT^-BirA* and TTBK2^fam1^-BirA*, and uncovered a strong enrichment of peroxisomal proteins in the TTBK2^fam1^-proximate data set. Using human telomerase-immortalized retinal pigment epithelial-1 cells (hTERT-RPE1), we then discovered and characterized localization of two distinct SCA11-associated proteins at peroxisomes (referred as TTBK2^fam1^ and TTBK2^fam2^ or TTBK2^SCA11^ proteins throughout the manuscript). Surprisingly, we also find that expression of SCA11-associated proteins interferes with peroxisome dynamics, reducing peroxisome numbers in the cytoplasm and at the ciliary base. We postulate that the observed defects caused by SCA11-associated proteins hamper the ability of peroxisomes to deliver cholesterol to the primary cilium leading to defects in SHH signaling activation.

## RESULTS

### TTBK2-truncated proteins associated with SCA11 are targeted to peroxisomes and decrease peroxisome cell abundance

*TTBK2*^fam1^ and *TTBK2*^fam2^ frameshift mutations result in truncated protein products that retain the kinase activity but do not localize to the centrosome and therefore cannot mediate cilium assembly (Bouskila et al., 2011; Goetz et al., 2012). These TTBK2^SCA11^ proteins do interfere with the function of wild-type TTBK2 in cilium assembly and trafficking, and in SHH signal transduction. However, they do not physically associate with TTBK2^WT^, and the cellular and molecular mechanisms of dominant interference that lead to neuronal dysfunction/loss in SCA11 remain unclear (Bowie et al., 2018).

To investigate the mechanisms by which TTBK2^SCA11^ might interfere with TTBK2^WT^ function in mediating ciliary assembly and trafficking, we performed biotin proximity labeling (Roux et al., 2012) to compare the interactomes of TTBK2^fam1^ and TTBK2^WT^. *Fam1* refers to a frameshift mutation in *TTBK2* initially identified in a British family pedigree affected with SCA11 (Houlden et al., 2007). We expressed tetracycline-inducible TTBK2^fam1^ and TTBK2^WT^ fused to BirA* labeling enzyme in Flp-In tREx HEK293 cells (Gupta et al., 2015). Following induction of the cells with tetracycline and treatment with biotin (**Fig S1A**), we harvested lysates, affinity purified using streptavidin beads, and subjected the purified proteins to quantitative mass spectrometry. Using this approach, we identified a total of 489 proteins with increased abundance in TTBK2^WT^ relative to TTBK2^fam1^ and 601 with reciprocally increased abundance in TTBK2^fam1^ compared with TTBK2^WT^ (**Table S1**). GO analysis of the proteins enriched in our TTBK2^fam1^ dataset included a large number of mitochondrial (84, FDR= 1.1x 10^-9^) and peroxisomal proteins (18, FDR= 1.0×10^-5^, **Fig S1B,C**), while those in the TTBK2 dataset were enriched for microtubule cytoskeleton (107, FDR= 2.26x 10^-26^) and centrosome proteins, consistent with our previous TTBK2 proximity proteome (Loukil et al., 2021).

To assess the sub-cellular localization of TTBK2^SCA11^, we generated RPE1 stable cell lines expressing two distinct *TTBK2* frameshift mutations associated with SCA11 (Houlden et al., 2007). In control TTBK2^FL^ cells, expressing full-length (FL) TTBK2, immunofluorescence and super-resolution microscopy analysis shows the characteristic localization wild-type TTBK2 at the distal appendages of the mother centriole (**Fig 1A**, arrowhead). Both transient and stable expression of TTBK2^fam1^ and TTBK2^fam2^ mutant proteins lack the well-known centriolar localization of TTBK2 and display instead a punctate cytoplasmic staining pattern (**Fig S2A, B**). Given that we identified many peroxisomal proteins among our TTBK2^fam1^ proximity data set (**Fig S1**), this led us to examine the co-localization of TTBK2^fam1^ and TTBK2^fam2^ proteins with peroxisomal markers. Both TTBK2^SCA11^ variants localize inside the peroxisomes labeled with the 70-kDa peroxisomal membrane protein (PMP70) (**Fig 1A**, insets) and the peroxisomal biogenesis factor 14 (PEX14) (**Fig S3A**). TTBK2^SCA11^ immunofluorescent signals closely overlaps with the signals of the peroxisomal β-oxidation enzyme Acyl-CoA Oxidase 1 (ACOX1), suggesting that these proteins are located within the lumen of peroxisomes (**Fig 1B**, insets). Immunofluorescent images for the ubiquitous peroxisomal membrane marker, PMP70, indicate that TTBK2^fam1^ and TTBK2^fam2^ cells display larger peroxisomes with altered morphology, and are also noticeably reduced in number when compared with TTBK2^FL^ control cells (**Fig 1C**). Quantification revealed that cells expressing TTBK2^fam1^ and TTBK2^fam2^ pathogenic proteins have nearly a 50% reduction in peroxisome number (mean of 166 vs 80 peroxisomes per cell) (**Fig 1D**). TTBK2^fam1^ and TTBK2^fam2^ cells also present a ∼2.5-fold increase in both PMP70 fluorescence intensity and peroxisome volume (**Fig 1E, F**). By contrast, no peroxisomal localization of TTBK2^FL^ was observed (insets, **Fig 1A, B**).

**Figure 1.**
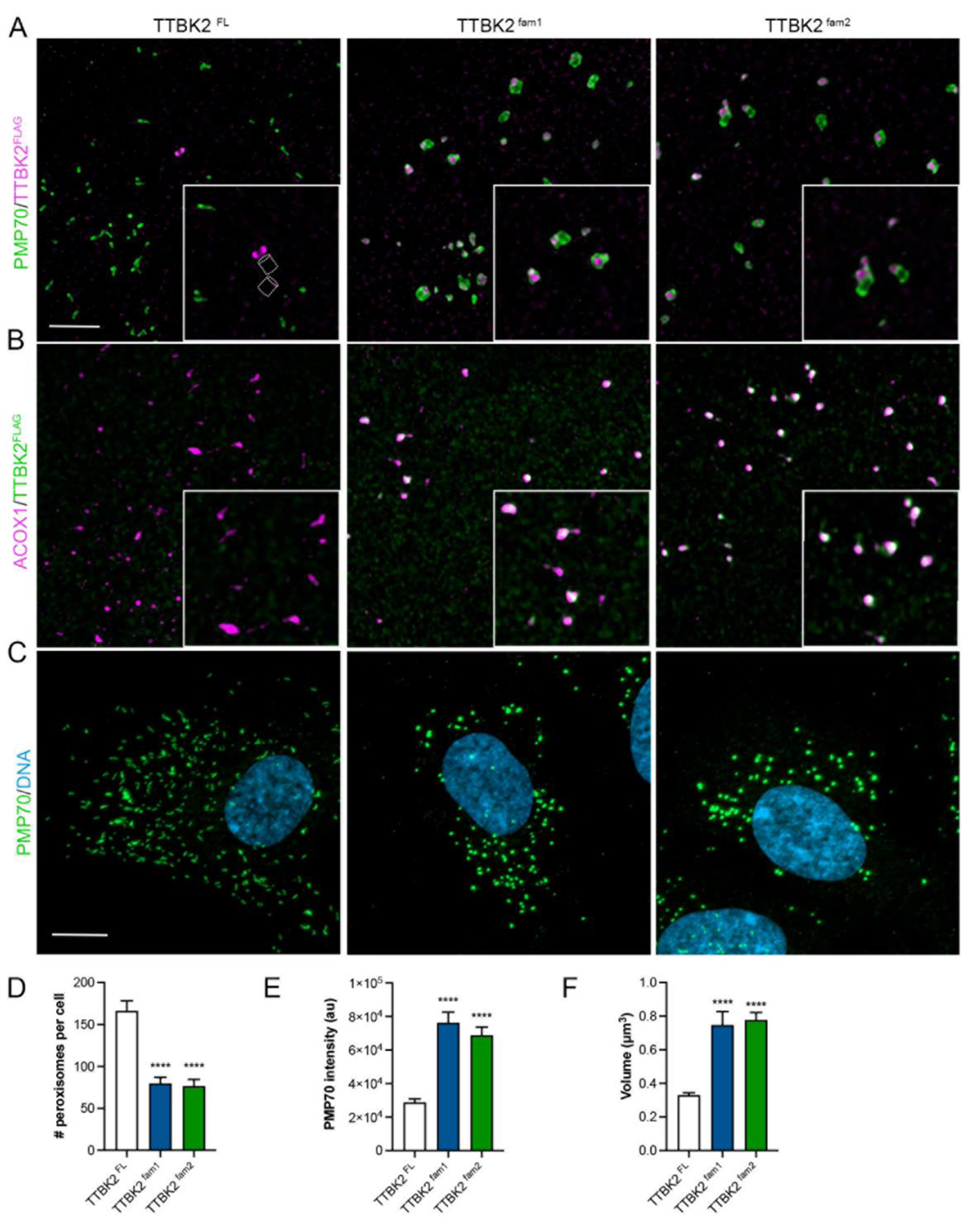
SCA11-causing frameshift mutations in *TTBK2* generate products which are trafficked to peroxisomes and caused peroxisome defects. (A) Super-resolution fluorescent images of stable RPE1 cell lines expressing full-length TTBK2 (TTBK2^FL^) or truncated SCA11-causing proteins (TTBK2^fam1^ and TTBK2^fam2^). Of note, all proteins were tagged with GFP-FLAG in their N-terminal domain. Cells were co-immunostained with antibodies against FLAG (magenta) and PMP70 (green) proteins or (B) FLAG (green) and ACOX1 (magenta) proteins. FLAG detects localization of GFP-FLAG-fusion proteins, and PMP70 and ACOX1 are membrane and matrix peroxisomal makers, respectively. Insets show the characteristic localization of wild-type TTBK2 protein at the centrioles in TBK2^FL^ cells (outlined cylinders depicts centrioles; see also **Figure S1A**). In stark comparison, TTBK2^fam1^ and TTBK2^fam2^ pathogenic proteins abnormally localize to peroxisomes. Bar, 2.5 μm. (C) Fluorescent images of TTBK2^FL^, TTBK2^fam1^ and TTBK2^fam2^ cells where peroxisomes are immunolabeled with PMP70 peroxisome marker (green) and DNA/nuclei (Hoechst 33342, blue). Bar, 10 μm. (D) Quantification of peroxisome cell number, (E) PMP70 signal intensity and (F) volume of individual peroxisomes per cell. Bars represent mean ± s.e.m.: ****P<0.0001: Kruskal-Wallis test with Dunn’s multiple comparisons test or one-way ANOVA with Tukey’s multiple comparisons test, n=2;15 cells per experiment.

### SCA11-associated variants contain a PTS1 and their kinase activity contributes to peroxisomal defects

Peroxisomes are single-bounded membrane organelles ubiquitously present in eukaryotic cells with important functions in metabolism, redox homeostasis, and human health (Steinberg et al., 2006; Walker et al., 2018). Most proteins targeted to the peroxisome have a consensus <u>p</u>eroxisomal target <u>s</u>ignal (PTS) in the extreme C-terminal or type 1 (PTS1) consisting of a tripeptide with a variable sequence (S/A/C), (R/K/H), (L/M) (Farre et al., 2019; Gould et al., 1989). Our analysis of the amino acid sequence of TTBK2^fam1^ and TTBK2^fam2^ proteins revealed an *in silico* predicted PTS1 (Emanuelsson et al., 2003) consisting in serine, glutamine and leucine (or SQL) tripeptide (**Fig 2A**). Similarly, analysis of the amino acid sequence of the other two frameshift mutations in *TTBK2* linked with SCA11 showed that the encoded protein products also contain SQL tripeptide in the C-terminal domain (**Fig S3C**). This suggests that all these genetic alterations in *TTBK2*, identified in different family pedigrees worldwide, share a common molecular pathogenic mechanism. We then assessed whether the SQL motif was responsible for abnormally targeting TTBK2^fam1^ and TTBK2^fam2^ proteins to peroxisomes. Previous evidence showed that the leucine (L) at −1 position from the PTS1 in NDR2 kinase sequence is essential to retain its peroxisomal localization (Abe et al., 2017). We therefore generated constructs to stably express TTBK2^fam1^ and TTBK2^fam2^ proteins lacking the terminal leucine (L) in RPE1 cells (TTBK2^fam1ΔL^ and TTBK2^fam2ΔL^ cell lines). In TTBK2^FL^ cell cultures, western blot analysis of GFP-fused cell fractions, a main band of ∼250 kDa was observed, which corresponds to TTBK2 full-length fusion protein. In cells expressing either TTBK2^fam1^, TTBK2^fam1ΔL^, TTBK2^fam2^, TTBK2^fam2ΔL^ protein, a band of ∼130 kDa was identified in all cell lines, indicating that truncated proteins associated with SCA11 are expressed despite leucine removal from the predicted PTS1 (**Fig 2B**). Strikingly, super resolution microscopy analysis in TTBK2^fam1ΔL^ and TTBK2^fam2ΔL^ proteins are no longer trafficked to peroxisomes (**Fig 2C**, insets). Moreover, in both cell lines the rounded and tubular morphology of peroxisomes is comparable to control cells (TTBK2^FL^, **Fig 1A**). We also found that in TTBK2^fam1ΔL^ and TTBK2^fam2ΔL^ cell cultures, peroxisome traits including number, PMP70 intensity, and volume are reestablished to TTBK2^FL^ control values (**Fig 2D-F**). This demonstrates that SQL tripeptide present in the sequence of SCA11-associated TTBK2 variants is a *bona fide* PTS1. Moreover, the abnormal trafficking of these proteins to peroxisome disrupts morphology of these microbodies and lead to a significant reduction in their number within the cell.

**Figure 2.**
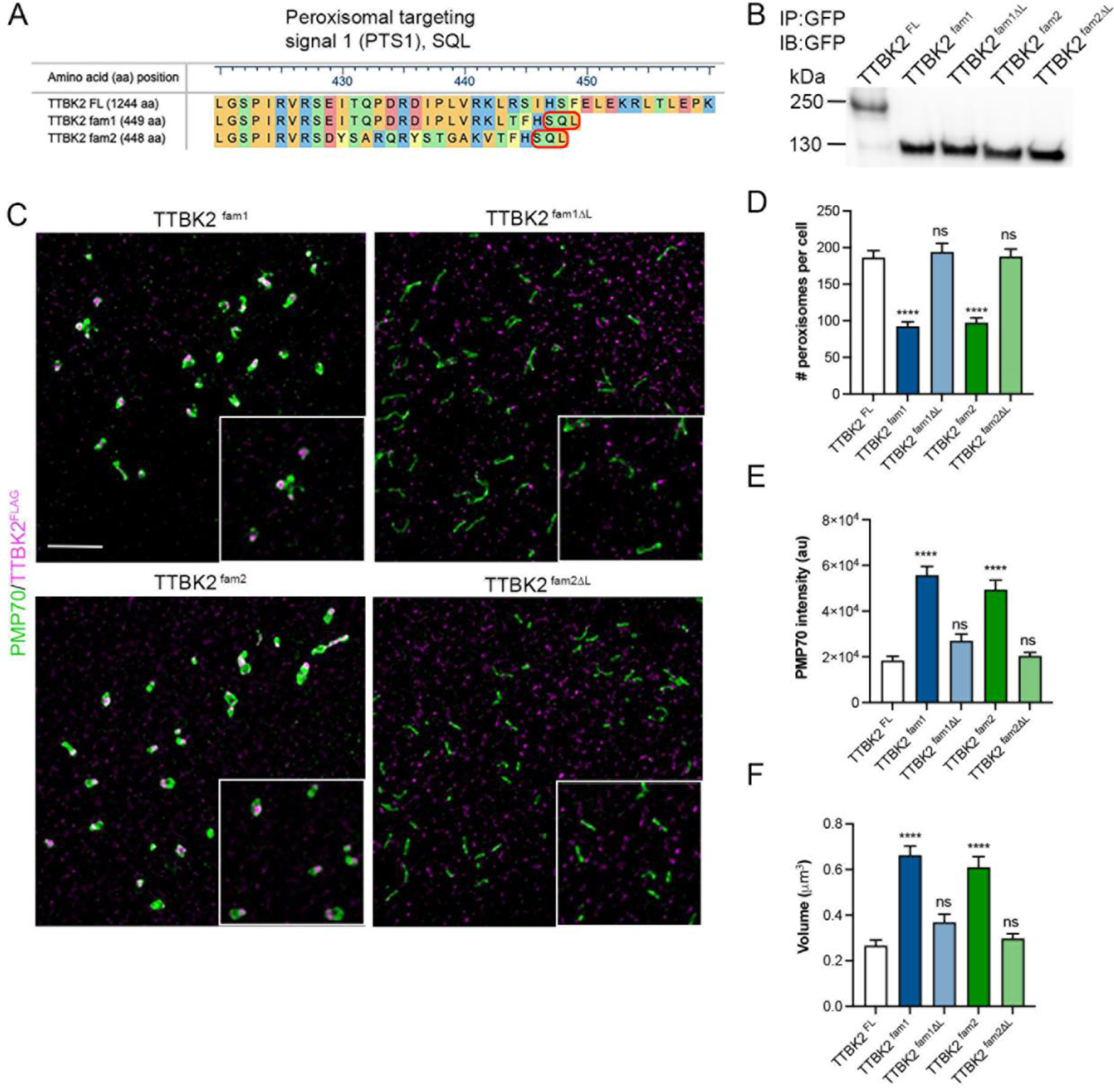
SCA11-causing frameshift mutations in *TTBK2* result in truncated proteins with a *bona fide* <u>p</u>eroxisomal targeting <u>s</u>ignal type 1 (or PTS1). (A) Scheme shows protein sequence alignment of wild-type TTBK2 (420–460 amino acids, aa) with the extreme C-terminal domain of TTBK2^fam1^ and TTBK2^fam2^ proteins. TTBK2^fam1^ and TTBK2^fam2^ are SCA11-causing truncated proteins of approximately 450 aa, where “Serine, glutamine and leucine” (or S, Q, L) is a predicted PTS1 result of the change in *TTBK2* reading frame. (B) Immunoprecipitation and immunoblotting analysis using GFP-Trap Agarose and a GFP antibody in cell lysates of TTBK2^FL^, TTBK2^fam1^, TTBK2^fam1ΔL^, TTBK2^fam2^ and TTBK2^fam2ΔL^ stable RPE1 cell lines. All lines express GFP/FLAG fusion proteins and TTBK2^fam1ΔL^ and TTBK2^fam2ΔL^ express TTBK2^fam1^ and TTBK2^fam2^ proteins lacking the leucine (L) from the predicted PTS1. (C) Super-resolution fluorescent images of TTBK2^fam1^, TTBK2^fam2^, TTBK2^fam1ΔL^ and TTBK2^fam2ΔL^ cells co-immunostained for FLAG (magenta) and PMP70 (green) as peroxisome membrane marker. Insets show loss of peroxisomal localization of TTBK2^fam1^ and TTBK2^fam2^ proteins after removal of the leucin (ΔL) from the predicted PTS1. Bar, 2.5 μm. (D) Quantification of peroxisome cell number, (E) PMP70 signal intensity and (F) volume of individual peroxisomes per cell. Bars represent mean ± s.e.m.: ns (not significant), ****P<0.0001: Kruskal-Wallis test with Dunn’s multiple comparisons test or one-way ANOVA with Tukey’s multiple comparisons test, n=2;15 cells per experiment.

SCA11-associated TTBK2 variants lack the C-terminal region of the protein but retain the N-terminal kinase domain. Several lines of evidence situate phosphorylation as an important regulatory mechanism of peroxisome dynamics (Oeljeklaus et al., 2016). This prompted us to determine the contribution of kinase activity in the peroxisomal phenotypes we observed in cells expressing the SCA11-associated variants. We generated and stably expressed kinase dead (D163A) analogs of fam1 and fam2 proteins in RPE1 cells (TTBK2^fam1KD^ and TTBK2^fam2KD^ cell lines), and observed that these proteins still localized at peroxisomes, as labeled with PMP70 (**Fig 3A**). Western blot analysis showed that TTBK2^fam1KD^ and TTBK2^fam2KD^ proteins are expressed at a slightly lower molecular weight than their kinase intact analogs (∼140 KDa), possibly indicative of autophosphorylation in the kinase active variants (**Fig 3B**). In cells expressing TTBK2^fam1KD^ and TTBK2^fam2KD^, peroxisome number was comparable to control values (mean, ∼166 vs 200 peroxisomes per cell) (**Fig 3C**). Furthermore, their expression does not alter PMP70 intensity or peroxisome volume when compared to TTBK2^FL^ control cells (**Fig 3D, E**). These results highlight kinase activity as an important effector of the peroxisomal alterations observed in cells expressing TTBK2^fam1^ and TTBK2^fam2^.

**Figure 3.**
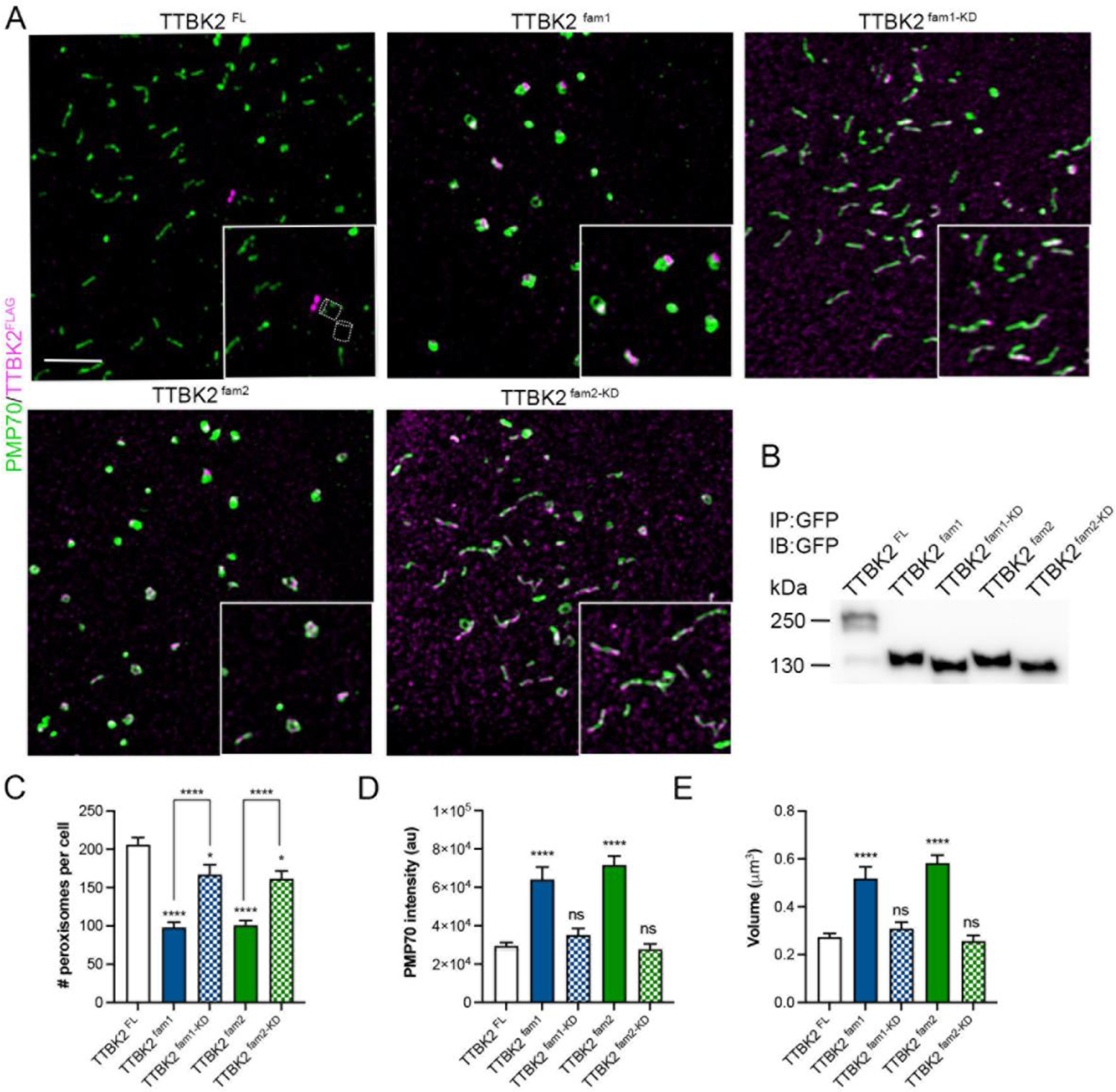
Kinase activity of SCA11-causing proteins mediates peroxisome defects. (A) Super-resolution fluorescent images of stable RPE1 cell lines expressing full-length TTBK2 (TTBK2^FL^), TTBK2^fam1^ and TTBK2^fam2^, or TTBK2^fam1^ and TTBK2^fam2^ kinase dead analogous (TTBK2^fam1KD^ and TTBK2^fam2KD^). All proteins were tagged with GFP-FLAG reporter. Cells were immunolabeled for FLAG (magenta) and PMP70 (green) as peroxisome. Insets show peroxisomal localization TTBK2^fam1KD^ and TTBK2^fam2KD^ and outlined cylinders represent centrioles. Bar, 2.5 μm. (B) Immunoprecipitation and immunoblotting analysis by using GFP-Trap Agarose and a GFP antibody in cell lysates of TTBK2^FL^, TTBK2^fam1^, TTBK2^fam1KD^, TTBK2^fam2^ and TTBK2^fam2KD^. (C) Quantification of peroxisome cell number, (D) PMP70 signal intensity and (E) volume of individual peroxisomes per cell. Bars represent mean ± s.e.m.: ns (not significant), * P<0.05, ****P<0.0001: Kruskal-Wallis test with Dunn’s multiple comparisons test or one-way ANOVA with Tukey’s multiple comparisons test, n=2;15 cells per experiment.

### SCA11-associated proteins impair peroxisome MFF/DRP1 fission pathway and peroxisome dynamics

One existing model of peroxisome biogenesis posits that these organelles are primarily generated by the division of existing peroxisomes (Farre et al., 2019). In mammals, this model requires membrane elongation, constriction and final scission mediated by the GTPase activity of Dynamin-related protein 1 (DRP1) (Williams et al., 2015). The mitochondrial fission factor (MFF), a tail-anchored membrane protein, acts as a major adaptor for the recruitment of DRP1 to the peroxisomal membrane (Gandre-Babbe and van der Bliek, 2008). Our data suggest that peroxisome growth and division are disrupted by the anomalous presence of TTBK2^fam1^ and TTBK2^fam2^ proteins in these organelles. Whereas no changes in DRP1 and MFF expression or phosphorylation in serine regulatory residues (Bindels et al., 2017) of these two proteins were found in TTBK2^fam1^ cell cultures (**Fig S4**), co-immunolabeling of MFF and PMP70 showed an abnormal accumulation of MFF at the peroxisomal membrane in TTBK2^fam1^ and TTBK2^fam2^ cells compared to cells expressing kinase dead analogs (**Fig 4A**, insets). Results showed a ∼2-fold increase of MFF signal intensity in cells expressing TTBK2^SCA11^ pathogenic variants (**Fig 4B**). Similar to MFF defects, a significant accumulation of DRP1 is also observed at peroxisomal membranes in TTBK2^SCA11^ cells (**Fig 4C, D**).

**Figure 4.**
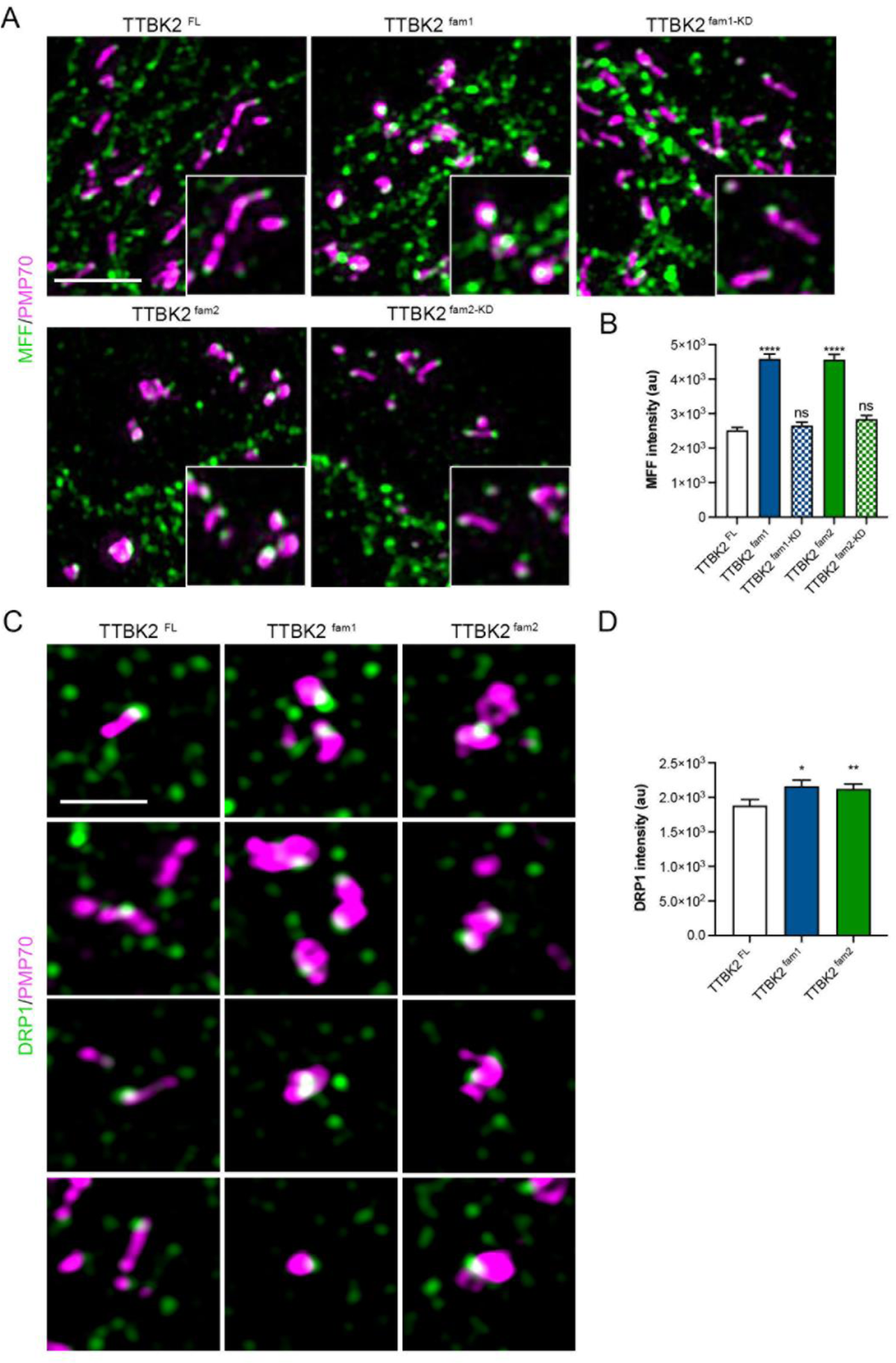
SCA11-causing proteins accumulate membrane fission components MFF/DRP1 at peroxisomal membranes. (A) Super-resolution fluorescent images of TTBK2^FL^, TTBK2^fam1^, TTBK2^fam2^, TTBK2^fam1KD^, TTBK2^fam2KD^ cells immunolabeled for PMP70 (magenta) and mitochondrial fission factor (MFF, green). Insets show MFF localization at peroxisomal membranes where noticeable abnormal accumulation is observed in TTBK2^fam1^ and TTBK2^fam2^ cells. Bar, 2.5 μm. (B) Quantification of MFF signal intensity at individual peroxisomes. (C) Insets of TTBK2^FL^, TTBK2^fam1^ and TTBK2^fam2^ cells immunolabeled for PMP70 (magenta) and dynamin-related protein 1 (DRP1, green), showing localization of DRP1 at the peroxisome membranes. Bar, 0.5 μm. (D) Quantification of DRP1 signal intensity at individual peroxisomes. Bars represent mean ± s.e.m.: ns (not significant), * P<0.05, **P<0.01, ****P<0.0001: Kruskal-Wallis test with Dunn’s multiple comparisons test, n=150 peroxisomes;10 cells for (B), n=200 peroxisomes;10 cells for (C).

To further examine potential defects in peroxisome fission, we carried out live cell imaging experiments in TTBK2^FL^ and TTBK2^fam1^ cells transiently expressing mScarlet protein fused with a PTS1 (**Fig 5A, B**). Collection of z-stacks images in living TTBK2^FL^ and TTBK2^fam1^ cells were obtained continuously with a time duration of 92.33 milliseconds per slice and result in 3D image stacks with approximately 164 frames (∼8min) for individual cells (**Video 1, 2**). Over the course of movies, from 3 to 6 fission events in peroxisomes were more frequently observed in TTBK2^FL^ cells in comparison to 0-2 events occurring in TTBK2^fam1^ cells (**Fig 5C**). A more complete visualization of the tracked peroxisomes in is shown in (**Fig S51, S52**). Interestingly, in TTBK2^FL^ cells, we frequently observed fusion of two fluorescent particles after they divide or fusion with other proximal fluorescent particle (**Fig 5A**, magenta arrows and F symbol, respectively). Similarly, to division, the frequency of fusion events is reduced in cells expressing TTBK2^fam1^ protein (**Fig 5D**). Notably, in TTBK2^fam1^ cells, peroxisomes failing to complete a division event form larger structures with various abnormal morphologies including long tubules and doughnut-like structures (**Fig 5E**). We conclude that SCA11-associated proteins trafficked to peroxisomes alter MFF/DRP1-dependent peroxisome fission pathway leading to a significant decrease in peroxisome cell number.

**Figure 5.**
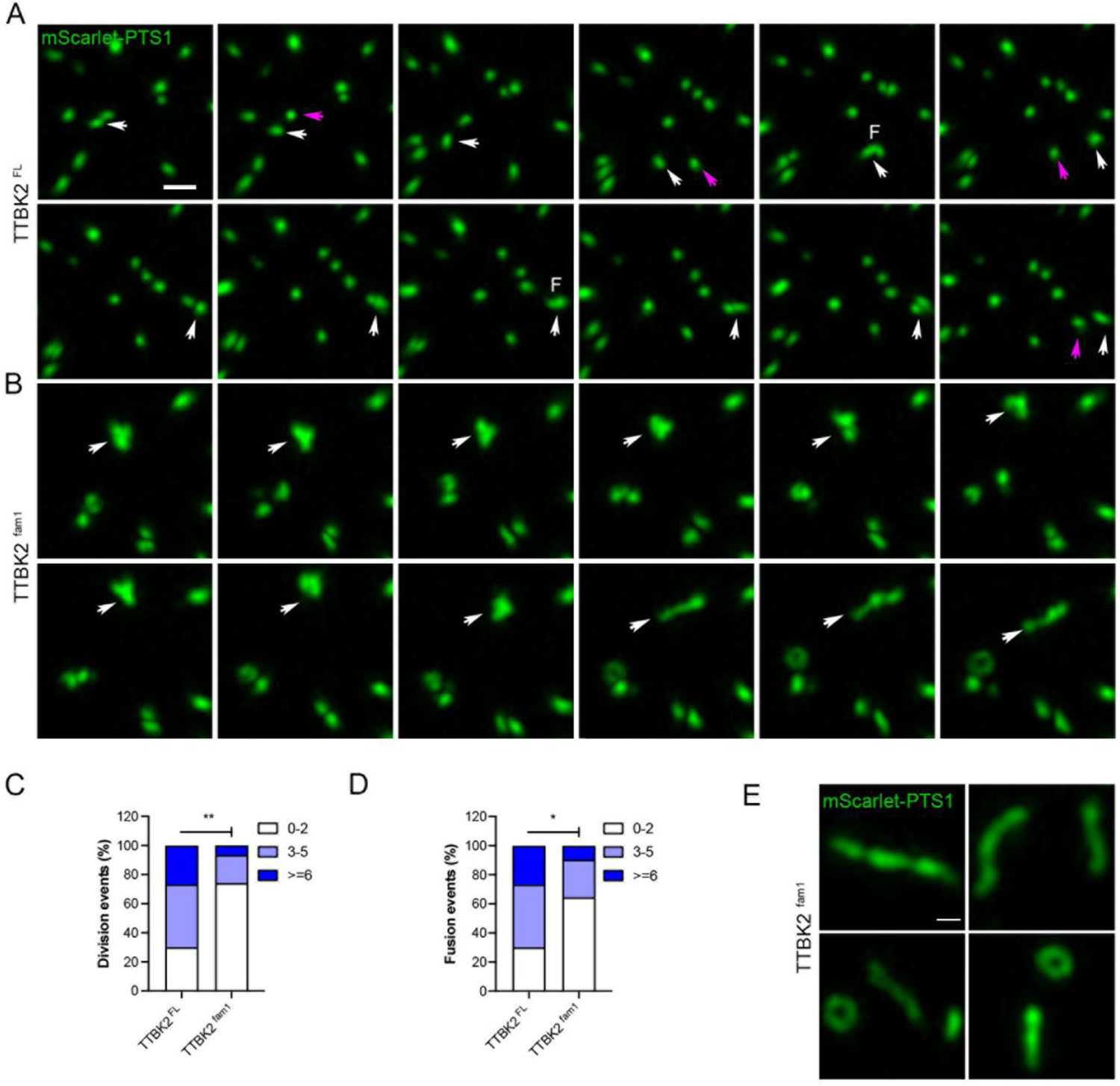
SCA11-causing proteins alter peroxisome dynamics. (A) Time-lapse analysis of peroxisome dynamics in living TTBK2^FL^ and (B) TTBK2^fam1^ cells transiently expressing mScarlet-PTS1 fluorescent protein. (A, B) Representative confocal images captured over the duration of the movies (∼8 min). Z-stack confocal images were recorded continuously with a frame time duration of 92.33 milliseconds resulting in 3D movies with a total of 164 frames for TTBK2^FL^ and TTBK2^fam1^ cells (**Video 1, 2**). (A, B) White arrows indicate the fluorescent particle analyzed overtime, magenta arrows and “F” character depicts division and fusion events, respectively. Bar, 10 μm. (C, D) Distribution of peroxisome division and fusion events represented as stack bars and expressed as percentages. *P<0.05, **P<0.01: Pearson’s chi-square (λ^2^) tests, n=30 peroxisomes; 2 cells per condition. (E) Fluorescent images of abnormal peroxisome morphologies observed in TTBK2^fam1^ cells. Bar, 0.5 μm.

### SCA11-causing proteins interfere with the localization of SHH ciliary components

In addition to the peroxisome defects here noted, SCA11-associated variants interfere with SHH signaling and cilia function in mice (Bowie et al., 2018). Moreover, peroxisomes deliver cholesterol to ciliary membranes (Miyamoto et al., 2020) where it plays a critical role in SHH signaling activation (Kinnebrew et al., 2021). Whereas no changes in cilia formation were detected in TTBK2^fam1^ and TTBK2^fam2^ cultured cells when compared to control cells (**Fig S8C**), a smaller fraction of TTBK2^SCA11^ ciliated cells displayed PMP70-positive vesicles near the ciliary base identified with γ-tubulin labeling (**Fig 6A**, insets). In TTBK2^FL^, TTBK2^fam1ΔL^ and TTBK2^fam2ΔL^ cells around 20% of ciliated cells have PMP70-positive vesicles near the ciliary base and this fraction is reduced by half in cells expressing TTBK2^fam1^ or TTBK2^fam2^ protein (**Fig 6B**). Similarly, another study reported approximately a frequency of 18% of cilia-bound peroxisome in wild-type RPE1 cells (Miyamoto et al., 2020).

**Figure 6.**
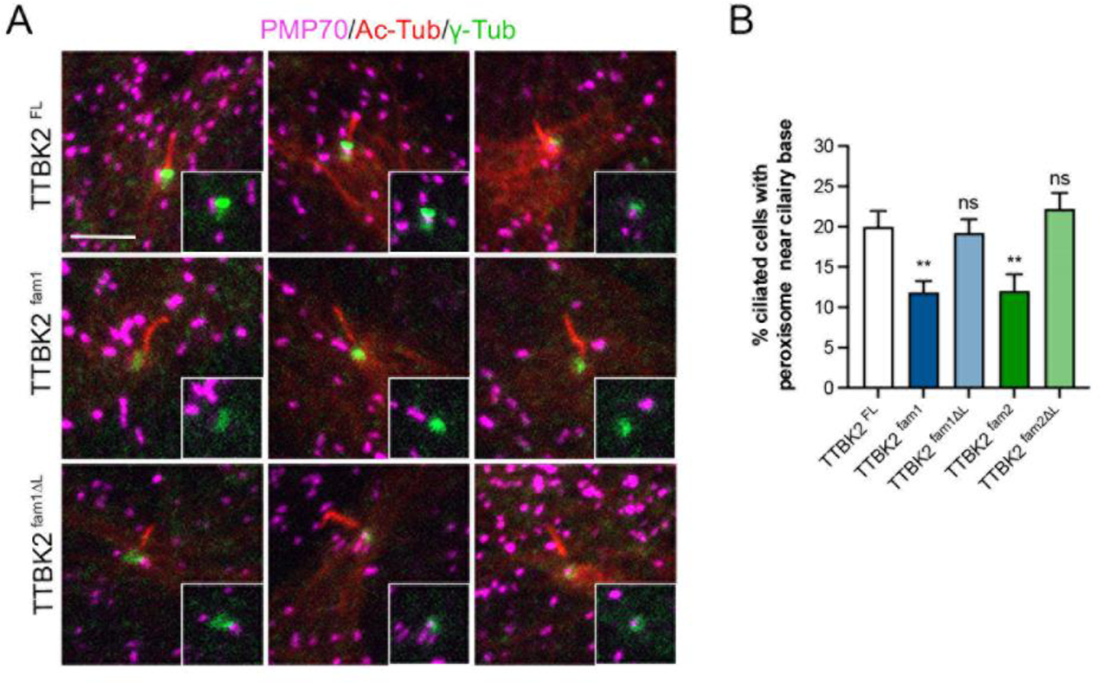
SCA11-causing proteins reduce peroxisomal interaction with cilia. (A) Fluorescent images of TTBK2^FL^, TTBK2^fam1^ and TTBK2^fam1ΔL^ ciliated cells immunostained for PMP70 (magenta), Acetylated α-Tubulin (Ac-Tub, red), γ-Tubulin (γ-Tub, green) as peroxisome, ciliary axoneme, basal body markers. For cilia induction cells were serum-deprived for 48 h. Insets show peroxisomes near the ciliary base, and a reduced interaction in these organelles is observed in TTBK2^fam1^. Bar, 5 μm. (B) Quantification of cilia-bound peroxisomes expressed as percentage in TTBK2^FL^, TTBK2^fam1^, TTBK2^fam1ΔL^, TTBK2^fam2^, TTBK2^fam2ΔL^. Bars represent mean ± s.e.m.: ns (not significant), *P<0.05, **P<0.01: Pearson’s chi-square (λ^2^) tests, n=2; 250 cells.

Based on these results, we hypothesized that ciliary cholesterol concentrations might be defective in TTBK2^fam1^ and TTBK2^fam2^ cells, where peroxisome abnormalities are observed. Unfortunately, we have been able to detect ciliary cholesterol in RPE1 cell lines by using Filipin III or mutant His_6_-tagged Perfringolysin (PFO*) fluorescently label probe (Kinnebrew et al., 2021) (**Fig S6B, C**), suggesting that ciliary membranes in RPE-1 cells contain very low concentrations of cholesterol, in agreement with a previous report (Breslow et al., 2013).

To overcome this challenge, we tested whether peroxisome defects in cells expressing SCA11-associated proteins could be interfering with SHH ciliary function attributable to cholesterol. We assessed ciliary accumulation of the G-protein-coupled receptor Smoothened (SMO), in TTBK2^FL^, TTBK2^fam1^ and TTBK2^fam1ΔL^ cell cultures. As expected, cells treated with SHH-N accumulate SMO along the ciliary membrane immunodetected with ARL13B protein (**Fig 7A**, Control, arrowheads). In contrast, cultures depleted of cholesterol using methyl-β-cyclodextrin (MβCD) in the presence of pravastatin (cholesterol biosynthesis inhibitor) and treated with SHH-N, ciliary SMO accumulation is inhibited (**Fig 7A**, MβCD). When cholesterol was extracted but exogenously supplemented in two forms (low-density lipoprotein, LDL-cholesterol, and water-soluble cholesterol), ciliary SMO accumulation is restored only in TTBK2^FL^ and TTBK2^fam1ΔL^ cells (**Fig 7A**, LDL and Chol^(+)^, arrowheads). In contrast, a lower frequency in SMO-positive cilia was observed in TTBK2^fam1^ cells (**Fig 7A**, LDL and Chol^(+)^, **Fig S7**). In agreement with these observations, in control conditions approximately 30% ciliated cells accumulate SMO at the ciliary membrane, this percentage decreased to ∼5% in cholesterol-depleted conditions in all cell cultures (**Fig 7B**). Notably, cholesterol supplementation in TTBK2^FL^ and TTBK2^fam1ΔL^ cells partially reestablished ciliary SMO accumulation (∼20%) but this restorative effect is reduced by half in TTBK2^fam1^ cell cultures (**Fig 7B, Fig S7**). Concomitant with the decrease in the proportion of SMO-positive cilia in TTBK2^fam1^ cells, a significant reduction in SMO ciliary intensity is also observed (**Fig 7C**). Interestingly, in TTBK2^fam1^ cell cultures, we also find a slight decreased in ARL13B ciliary levels (**Fig S7B**). In summary this data highlights the importance of peroxisomes and cholesterol in the trafficking of SHH signaling components to cilia and suggest that defects in the SHH pathway could a contributing factor of SCA11 neuropathology.

**Figure 7.**
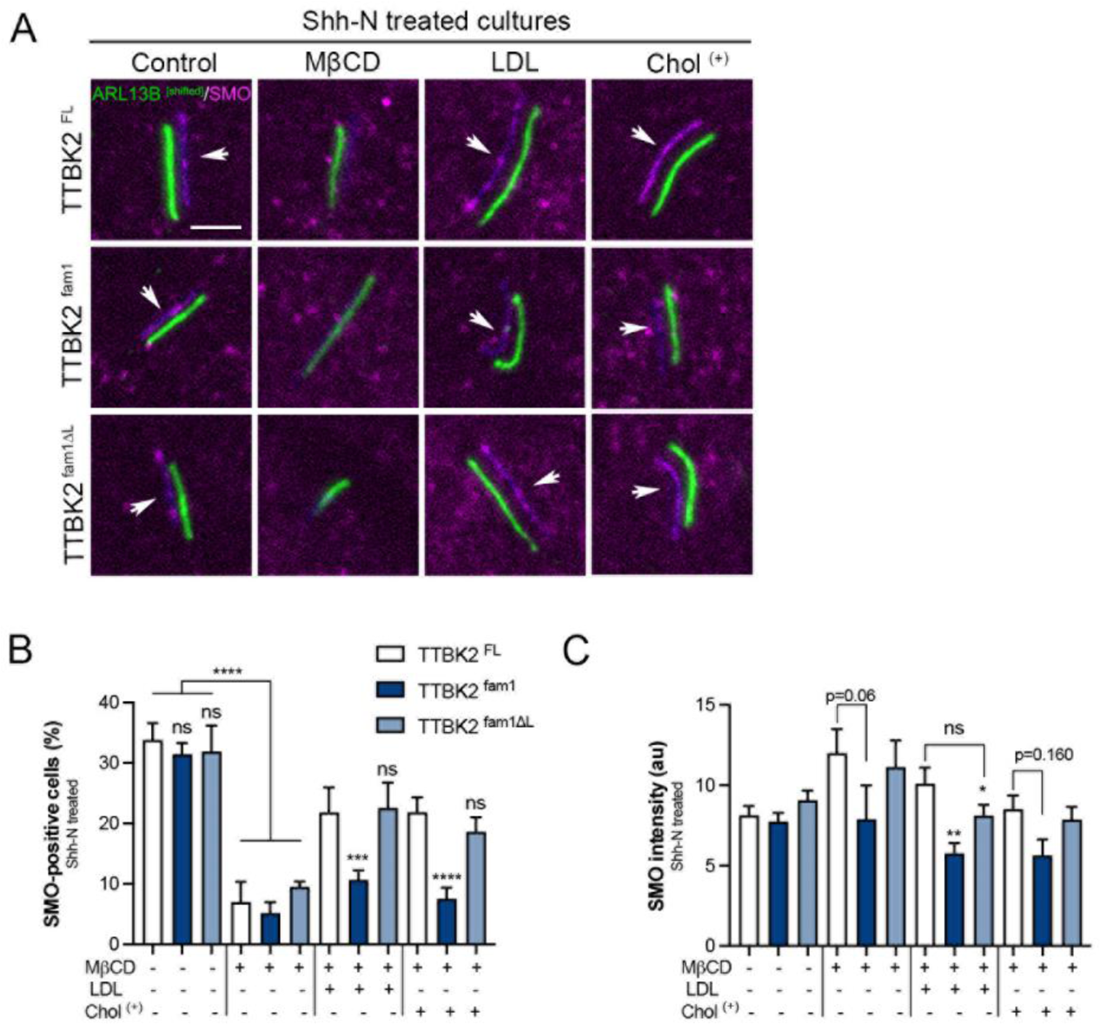
SCA11-causing proteins reduce trafficking of ciliary SMO. (A) Immunofluorescent images of cilia in TTBK2^FL^, TTBK2^fam1^, TTBK2^fam1ΔL^ cells treated with 50 nM of SHH-N for 24 h (Control) and immunolabeled for SMO (magenta) and ARL13B (green) ciliary membrane markers. Parallel cultures were depleted of cholesterol using Methyl-β-cyclodextrin (MβCD) and either supplemented with low*-*density lipoprotein cholesterol (LDL) or water-soluble cholesterol (Chol^+^) in the presence of 50 nM of SHH-N and 40 μM of cholesterol biosynthesis inhibitor pravastatin for 24h. Arrows point out SMO localization along the ciliary membrane upon SHH-N stimulation. Bar, 2.5 μm. (B) Quantification of proportions of SMO-positive cells expressed as percentage and (B) SMO ciliary intensity in TTBK2^FL^, TTBK2^fam1^, TTBK2^fam1ΔL^ cell cultures with the indicated treatments. For (B) see also Fig 8S. Bars represent mean ± s.e.m.: ns (not significant), *P<0.05, **P<0.01, ***P<0.001: Pearson’s chi-square (λ^2^) tests for (B), Kruskal-Wallis test with Dunn’s multiple comparisons test for (C), n=2; 250 cells.

## DISCUSSION

The primary cilium is an incomplete membrane-bounded cellular projection that extends into the extracellular environment and is present as a single structure in most vertebrate cell-types. By contrast, peroxisomes are single-membrane-bounded organelles distributed throughout the cytoplasm. A large number of these microbodies (10^2^-10^3^ per cell) are present in human cells, with numbers varying by cell-type. Cilia and peroxisomes nonetheless share several key characteristics important for cellular function: 1) they exhibit dynamic and highly regulated trafficking of cytoplasmic proteins to these subcellular compartments, 2) they have important functions in sensing and responding to environmental cues activating cell signaling pathways, and 3) dysfunction of each of these organelles causes hereditary developmental syndromes: “ciliopathies” and peroxisome biogenesis disorders (PBDs) respectively (Reiter and Leroux, 2017; Steinberg et al., 2006). These groups of illnesses often result in early lethality or poor quality of life, and are characterized by systemic organ failure, including neurological abnormalities. To date, mutations in more than 70 genes have been reported to cause a spectrum of ciliopathies including Joubert syndrome, Meckel syndrome, and nephronophthisis (NPHP) (Van De Weghe et al., 2022). These gene mutations impair localization of the protein products to the cilia/centrosome compartment and result in defects in cilia formation and function. In addition, two independent studies have evidenced the participation of peroxisome-localizing proteins in cilia formation (Abe et al., 2017; Maharjan et al., 2020). Notably, a recent work described a peroxisome-associated protein complex important for supplying cholesterol to primary cilia membrane (Miyamoto et al., 2020), which is in turn important for activation of SHH signaling pathway (Radhakrishnan et al., 2020). In this study, we reveal how pathogenic mutations in the ciliary gene *TTBK2* disrupt cilia-dependent SHH signaling, in part, by causing defects in peroxisome dynamics. Our results point to a possible contribution of dysfunctional peroxisome-cilia interaction in spinocerebellar ataxia type 11 (SCA11) pathogenesis.

Frameshift mutations in *TTBK2* cause the hereditary, autosomal dominant neurodegenerative disorder SCA11 (Houlden et al., 2007). In SCA11-affected individuals, head MRI and postmortem tissue analysis indicate cerebellar atrophy and loss of Purkinje cells in the cerebellum. However, the molecular and cellular mechanisms associated with SCA11 pathology are not well established. Here, we discovered that two distinct SCA11-causing proteins (TTBK2^fam1^ and TTBK2^fam2^) are imported into the lumen of peroxisomes. The abnormal localization of these truncated proteins to peroxisomes is mediated by a <u>p</u>eroxisomal target <u>s</u>ignal type 1 (PTS1) (**Fig 1, 2**). Additional *TTBK2* frameshift mutations identified in SCA11-affected families are also predicted to produce truncated protein products that possess the PTS1 here characterized (**Fig S3C**). This points to a probable common pathogenic pathway in SCA11 involving peroxisome-dependent cellular processes. We find that translocation of TTBK2^fam1^ and TTBK2^fam2^ proteins to peroxisomes alters peroxisome growth and formation without perturbing the localization of peroxisome membrane proteins including, PMP70 and PEX14 (**Fig 1, S3A**). We also demonstrate that kinase activity of TTBK2^SCA11^ truncated proteins is necessary to trigger these peroxisomal defects (**Fig 3**). Although peroxisome abundance is decreased in cells expressing the truncated proteins, the remaining organelles are capable of transporting ACOX1 and Catalase, two major peroxisome hydrogen peroxide (H_2_O_2_)-metabolizing enzymes (**Fig 1B, S3B**). It is estimated that peroxisomes can account for ∼35% of H_2_O_2_ production in mammalian tissues (De Duve and Baudhuin, 1966). Strikingly, we do not detect significant changes in the level of H_2_O_2_ in TTBK2^fam1^ cells under normal culture conditions or upon menadione treatment (**Fig S8**), indicating that TTBK2^SCA11^ proteins do not alter the overall cellular metabolism of H_2_O_2_.

Phosphorylation of peroxisome-localizing proteins including MFF and DRP1 at multiple serine residues is reported to regulate peroxisome fission (Oeljeklaus et al., 2016). In cells expressing TTBK2^fam1^ protein, our analysis of regulatory residues of MFF (Ser147) and DRP1 (Ser616 and 637) indicate no detectable changes in serine phosphorylation (**Fig S4**). Future phospho-proteomic studies will determine whether alternative phosphorylation sites in MFF/DRP1 or other fission machinery components explain the peroxisomal defects we identified here. Nevertheless, cells expressing SCA11-associated proteins exhibit an accumulation in MFF and DRP1 at peroxisomes (**Fig 4**). Consistent with the decreased peroxisome abundance observed in fixed-cells expressing SCA11-causing mutations (**Fig 1**), cells expressing TTBK2^fam1^ protein also display fewer division events observed by live cell imaging analysis (**Fig 5**). MFF facilitates the recruitment of DRP1 to catalyze membrane fission in mitochondria (Gandre-Babbe and van der Bliek, 2008). However, TTBK2^SCA11^ truncated proteins do not localize to mitochondria or aberrantly accumulate MFF at the mitochondrion membrane in RPE1 cells (**Fig S9**), suggesting that mitochondria dynamics is not directly perturbed. Indeed, when disturbances in the dynamics of both peroxisome and mitochondria occur due to *MFF* pathogenic mutations, the affected individuals present clinical phenotypes associated with Leigh-like encephalopathy rather than SCA11 (Koch et al., 2016; Shamseldin et al., 2012). Therefore, our findings indicate that translocation of SCA11-causing proteins primarily affect membrane remodeling events during peroxisome growth and division by mechanisms entailing MFF-dependent peroxisome fission pathway.

Peroxisomes participate in intracellular distribution of cholesterol between organelles (Chu et al., 2015; Miyamoto et al., 2020). In Zellweger syndrome (ZS), a severe PBD characterized by multi-organ dysfunction, loss of functional peroxisomes results in massive intracellular accumulation of cholesterol (Santos et al., 1988; Small et al., 1988). Despite this intracellular accumulation of cholesterol, ZS patient-derived fibroblasts have a reduction in both ciliary membrane cholesterol and ciliary localization of SMO upon SHH signaling activation (Miyamoto et al., 2020). While no noticeable accumulation of intracellular cholesterol is found in cells expressing TTBK2^SCA11^ proteins (**Fig S6A**), ciliated cells exhibit fewer peroxisomes interacting with the ciliary base (**Fig 6**), suggesting defects in peroxisome subcellular distribution. In addition, upon cholesterol depletion and re-addition of exogenous cholesterol, cells expressing TTBK2^fam1^ protein exhibit a significant reduction in SMO-positive cilia and SMO intensity along the ciliary membrane upon treatment with SHH-N (**Fig 7**). These findings indicate that SCA11-associated proteins hamper the ability of peroxisomes to deliver ciliary cholesterol (Miyamoto et al., 2020) and contribute to explain their interference in cilia formation and ciliary molecular composition by unaltering the function of wild-type TTBK2 (Bowie et al., 2018).

In this study, we propose novel molecular mechanisms associated with SCA11 neuropathology and highlight the importance of the peroxisome-primary cilium intracellular communication for cellular homeostasis. The current work also provides a solid framework to further study this inter-organelle interaction in brain physiology with particular importance in the maintenance of Purkinje neurons in the cerebellum.

## MATERIALS AND METHODS

### BioID

Flp-IN tREx 293T cells stably expressing tetracycline-inducible GFP-, TTBK2^WT^, or TTBK2^fam1^-BirA were treated with 1µg/ml of tetracycline for 48 h to induce protein expression. Cells were then treated with 50µM of biotin and lysed 24 h later in 50mM Tris pH 7.5, 100mM NaCl, 0.5% NP-40, and 10% glycerol. Lysates were sonicated, cleared and incubated with Streptavidin-Sepharose beads. Biotinylation was validated by western blot, and samples were sent off to the Duke University Proteomics and Metabolomics Core Facility for mass spectrometry processing. Analysis of peptide abundance was performed as previously described (Loukil et al., 2021).

### Cell culture and retroviral transduction

Telomerase-immortalized human retinal pigment epithelial cells (hTERT RPE-1 or RPE1 cells) were purchased from ATCC. RPE1 cells were cultured in high-glucose DMEM/F-12 supplemented with 10% (vol/vol) FBS, 100 units/ml of penicillin and 100 µg/ml streptomycin at 37°C in a humidified 5% CO_2_ incubator (all reagents from Thermo Fisher Scientific). Stable RPE1 cells were generated by retroviral transduction. For retroviral production, retroviral vector containing TTBK2 and FLAG/eGFP sequences and envelop vector (Goetz et al., 2012) were transfected into Phoenix cells plated 100-mm dishes using Lipofectamine 2000 (Invitrogen). Fresh DMEM medium supplemented with 10% (vol/vol) FBS was changed 4 h post-transfection, and 2 days after, viral supernatants were collected and centrifuged at 10,000g for 5 min and filtered through 0.45-μm filters. RPE1 cells plated on 30-mm dishes (∼70% confluence) were infected with retroviruses (3ml of RPE1 growth media: 2ml of supernatant; supplemented with 2 µg/ml of polybrene) for 24 h and selected with 0.5 mg/ml of Geneticin/G418 Sulfate (Gibco) for at least one week. Stable RPE1 cells were cultured in high-glucose DMEM/F-12 supplemented with 10% (vol/vol) FBS and 0.5 mg/ml of Geneticin at 37°C in a humidified 5% CO2 incubator (all reagents from Thermo Fisher Scientific). Of note, all RPE1 stable cell lines here developed express TTBK2 proteins tagged in their N-terminal domain with FLAG/eGFP reporter. For simplification proposes, FLAG/eGFP labeling is omitted throughout the manuscript/figures when referring to stable expression of fusion proteins. *Fam1* and *fam2* refers to TTBK2 mutations initially identified in two family pedigrees affected with SCA11 (c.1329dupA and c.1284_1285delAG, DNA nucleotide change, respectively) (Houlden et al., 2007). For peroxisome analyses ∼4×10^4^ cells were seeded per well in 24-well plates and maintained in culture for at least 24 h.

### Constructs

All TTBK2 mutations were generated by PCR-based site-directed mutagenesis by using QuickChange Lighting Site-Directed-Mutagenesis kit (Agilent) using as a template pENTR/D-TOPO (Thermo Fisher Scientific) vector containing cDNA of human TTBK2. Sanger sequencing method was used for confirming TTBK2 mutations and primers were designed using QuikChange Primer Design Program (Agilent) (**Table S2**). TTBK2 inserts were cloned into retroviral expression vector pQCXIN (Clonetech) by carrying out Gateway cloning and for retroviral production pVSV-G pantropic envelope vector was used (Goetz et al., 2012). pmScarlet_peroxisome_C1 was used for in vivo visualization of peroxisomes and was a gift from Dorus Gadella (Addgene, 85063).

### Antibodies

For immunostaining, rabbit anti-PMP70 (Invitrogen, PA1-650, 1:1,000), mouse anti-FLAG (Sigma-Aldrich, F1804, 1:2,000), anti-ACOX1 (ABclonal, A8091, 1:1,000), rabbit anti-MFF (Proteintech, 17090-1-AP, 1:2,000), mouse anti-PMP70 (Sigma-Aldrich, SAB4200181, 1:1000), rabbit anti-DRP1 (Abcam, ab219596, 1:1000), mouse anti-γ-Tubulin (Sigma-Aldrich, T6567, 1:2000), mouse anti-Acetylated-Tubulin (Sigma-Aldrich, T6793, 1:2000), rabbit anti-Arl13b (Proteintech, 17711-11-AP, 1:2000), mouse anti-SMO (Santa Cruz, sc-166685, 1:500). Specific secondary fluorescent antibodies were used, all conjugated to Alexa flour dyes (Invitrogen, 1:1000). For western blot analysis, rabbit anti-GFP (Invitrogen, A-11122, 1:1,000). Anti-rabbit horseradish peroxidase (HRP)-conjugated secondary antibody was used at a 1:10,000 dilution (Jackson ImmunoResearch, 111-035-1400).

### Immunoprecipitation and Western blot analysis

Confluent monolayers plated in 10-mm dishes were washed with cold PBS and lysed on ice with NP40 lysis buffer (20 mM Tris–HCl, pH 8.0, 137 mM NaCl, 2 mM EDTA, and 1.0% NP-40, 5.0% Glycerol, containing a protease inhibitor cocktail (Roche) and 25 mM β-Glycerophosphate) for 30 min. Cell lysates were centrifuged 17,000x g for 10 min at 4°C and supernatants were incubated with ChomoTek GFP-Trap Agarose beads (Proteintech) for 1 h at 4°C followed by bead sedimentation and washing according to manufacturer’s instructions. Immunocomplexes were dissociated from beads with 2x Laemmi Sample buffer (Bio-Rad) and supernatant was subject to SDS-PAGE/Western blot analysis. Samples were resolved by 10% TGX stain-free polyacrylamide gels (Bio-Rad) and transferred to 0.45-μm PDVDF membranes (Thermo Fisher Scientific). Membranes were incubated in blocking buffer [5% (w/v) nonfat dry milk in Tris-buffered saline with 0.1% Tween (TBST)] for 1 h at room temperature followed by overnight incubation of GFP primary antibody (diluted in blocking buffer) at 4°C. Membranes were incubated for 1 h at room temperature with HRP-conjugated secondary antibody in TBST. Chemiluminescent detection was performed using the SuperSignal kit (Thermo Fisher Scientific) and quantified using a ChemiDoc MP imaging system with Image Lab 6.1 software (Bio-Rad).

### Immunofluorescence

Stable RPE1 cells grown onto glass coverslips during the indicated times were fixed with 4% PFA solution in PBS (Santa Cruz) and permeabilized with 0.20% Triton X-100 in PBS for 15 min. Then cells were incubating with blocking buffer [1% bovine serum albumin (BSA) and 5% goat serum in PBS] for 1 h at room temperature (RT) and stained using proper primary and fluorescent-dye-conjugated secondary antibodies (Alexa Fluor dyes, Invitrogen). Incubations of primary antibodies were performed overnight at 4°C and for fluorescently conjugated secondary antibodies 1h at RT. After four washes with PBS, coverslips were mounted using ProLong Diamond antifade mounting medium (Thermo Fisher Scientific). Widefield fluorescence microscopy was performed on an AxioObserver microscope (Zeiss) equipped with an HXP lamp (Zeiss), an AxioCam 506 mono camera (Zeiss), a Plan-Apochromat 63×/1.4N.A. oil objective (Zeiss). Image stacks were taken with a z distance of 0.25 µM and acquired in ZEN 3.4 software (Zeiss) using ApoTome.2 grid (Zeiss). Light intensity was set to 80% to minimize exposure times (≤ 50ms).

### Lattice Structured Illumination (SIM) Microscopy

Lattice-SIM images were acquired on a Zeiss Elyra 7 AxioObserver microscope equipped with an Plan-Apochromat 63×/1.4 Oil DIC M27 objective and two pco.edge sCMOS (version 4.2 CL HS) cameras which were aligned at the start of the experiment daily. The system contained 405 nm and 642 nm diode, and 488 nm and 561 nm OPSL lasers. For each focal plane 13 phase images were acquired. Exposure time and laser power were balanced for each fluorescence channel individually to minimize bleaching and set to values between 20-50 ms and 1-1.5 %, respectively. Alexa 488, 568 and 647 were excited using LBF 405/488/561/642 filter set and the emission was captured through an SBP 490–560 + LP 640 emission filter. SIM grid size was 36.5µm and SIM image processing and reconstruction was performed with the SIM processing tool of the ZEN 3.0 SR (black) software (Zeiss). Stacks were processing using 3D mode with default settings and channel alignment was apply to output images.

### SHH activation and cholesterol supplementation

Cells (∼7.5 ×10^4^) were seeded in wells of 24-well plates and maintained in culture for 30 h to reach confluence. Cells were then incubated for 24 h in serum-free DMEM/F-12 to induce cilia formation. Followed by 24 h treatment with 50 nM of Shh-N (R&D Systems) diluted in serum-free DMEM/F-12. For cholesterol supplementation, parallel RPE1 cell cultures, after cilia induction, were treated for 45 min with 1.5% methyl-β-cyclodextrin (MβCD, Sigma) in DMEM/F-12 to deplete cellular cholesterol. After extensive washing, cholesterol was delivered by incubating cells for 1 h with 100 μM of water-soluble methyl-β-cyclodextrin–cholesterol complex (Sigma-Aldrich) in DMEM/F-12. For LDL cholesterol complementation, cells were supplemented 0.034 mg/mL of LDL (MilliporeSigma) in DMEM/F-12, corresponding approximately to the amount of cholesterol supplemented with water-soluble methyl-β-cyclodextrin–cholesterol complex. Cholesterol depletion and supplementation assays were performed in the presence of both 40 μM pravastatin (Sigma-Aldrich) and 50 nM of Shh-N for 24 h. All samples were then processed for immunofluorescence to detect ciliary membrane components SMO and Arl13b.

### Live cell imaging

Stable RPE1 cells (∼0.5×10^6^) were electroporated with pmScarlet_peroxisome_C1 (or mScarlet-PTS1) (250 ng) using an Amaxa Nucleofector II (Lonza) in accordance with the manufacturer’s instructions using Ingenio electroporation solution (Mirus). Equal volumes of cell suspension were divided and plated in 3 wells of 8-well dishes (µ-Slide 8 well high glass bottom, Ibidi) and cells were maintained in culture for 15 h. Before live imaging, cells were washed three times and transferred to DMEM/F-12 with no phenol red (Thermo Fisher Scientific) supplemented with 10% (vol/vol) FBS. Airyscan images were acquired in the Multiplex SR-4Y mode on a Zeiss LSM 980 microscope equipped with an Airyscan 2 detector using a Plan-Apochromat 63×/1.4N.A. oil DIC M27 objective. LSM980 microscope was also equipped with an integrated incubation system and CO2-Cover for heating insert P, and cells were imaging at 37°C with 5 % CO2 conditions. Excitation of mScarlet protein was carried out with a 561 nm argon laser line (0.1%) and the emission was captured through an GaAsP-PMT Array Airyscan detector (570–620, detector gain 850). Pixel size in Airyscan high resolution acquisition was applied automatically in ZEN 3.5 software (Zeiss). The Pixel Dwell was 0.38 µsec. Z-Stack scans were performed at 0.17 µm intervals and time-lapse imaging was recorded for approximately 8 min. Images were deconvolved with the deconvolution tool of the ZEN Blue software (Zeiss). The acquired time series was processed through Airyscan Processing using Weiner Filter strength setting of 4.9.

### Image analysis

Peroxisome number, volume and PMP70 fluorescent intensity of individual peroxisomes were determined in widefield fluorescence Z-stack images of cells immunolabeled for PMP70. All these parameters were quantified by manually tracking individual cells in sub-confluent cell monolayers and by using 3D particle analysis (Bolte and Cordelieres, 2006) using a similar threshold for all images (ImageJ, NIH). For quantification of MFF/DRP1 signal intensity at peroxisomes, maximum intensity projections of SIM images and ImageJ were used. In the PMP70 fluorescent channel, a circle with a diameter of ∼0.35 µm was drawn in randomly selected peroxisomes and saved on the Region of Interest (ROI) manager. Then, in MFF/DRP1 and PMP70 merged channel images, ROI was corrected as necessary to enclose both PMP70 and MFF/DRP1 fluorescent signals. The sum of all the pixel intensities in the ROI was obtained for individual peroxisomes in MFF/DRP1 single channel images. Peroxisomes near the ciliary base were identified by PMP70-positive vesicle staining within an approximate radius of 1 µm from γ-tubulin staining using maximum intensity projections. To compare the intensity of ciliary SMO, integrated density values were determined for individual cilia by using a mask of Arl13b ciliary immunostaining. Arl13b-masks were generated by adjusting the threshold of maximum intensity projections for Arl13b channel and by adding ROI (individual cilia) using the wand-tracing tool. Percentage of cells with SMO-positive cilia were calculated manually considered as total the number of cilia identified by Arl13b labeling. Quantification of the number of division and fusion events in peroxisomes expressing mScarlet-PTS1 protein in vivo was performed in Imaris software (v.9.9.1). Time-lapse image analysis for randomly selected fluorescent particles was performed by observing their behavior in each time point (∼164 frames or time points per cell). Division was considered when no signal of mScarlet was detected between time points. For fusion, 3D images were rotated in each time to take into consideration z-dimension motion and discard overlapping of particles in 2D views. Time-lapse files were subject to bleach correction prior analysis in Imaris using ImageJ (NIH).

### Statistical analysis

For all data, we first assessed whether data were normally distributed. Depending on the distribution, data were analyzed by either ordinary One-way ANOVA with Tukey’s post hoc tests or Kruskal-Wallis with Dunn’s multiple comparisons. Statistical significance was set to P<0.05. P values and tests used are indicated in the corresponding figure legend and (n) denotes independent experiments, unless indicated otherwise. GraphPad Prism 9 (v.9.4.1) was used for create graphs and statistical analyses. Error bars indicate SEM. Figure preparation including display of representative fluorescent images and graphs was performed with Adobe Photoshop software (v.23.5.0).

## Acknowledgements

Thanks James Shaw and Garrick Koermer (Carl Zeiss Microscopy, LLC) for facilitating the use of a Zeiss LSM 980 microscope and valuable suggestions for live cell imaging experiments. Thanks Maia Kinnebrew and the Rohatgi lab (Stanford University) for providing PFO* fluorescent probe. We acknowledge the Duke University Light Microscopy (Lisa Cameron and Benjamin Carlson) (Grant Number: 1 S10 OD028703-01 for Elyra7 use) and Proteomics and Metobolomics core facilities. This work was supported by R01HD099784 and R03TR003392 to SCG. COI: SCG has consulted for Arvinas, Inc.

## Author contributions

JME and SCG conceptualization. JME designed and performed most of the experiments, including Site-Directed-Mutagenesis, development of stable cell lines, Lattice Structured Illumination (SIM) Microscopy imaging and live cell imaging experiments. AN performed and analyzed biotin proximity labeling experiment. JME and SCG, formal analysis. JME wrote the original draft and prepared figures. JME, AN, SCG, writing and review. SCG, project administration and funding acquisition.

---

**Figure S1.**
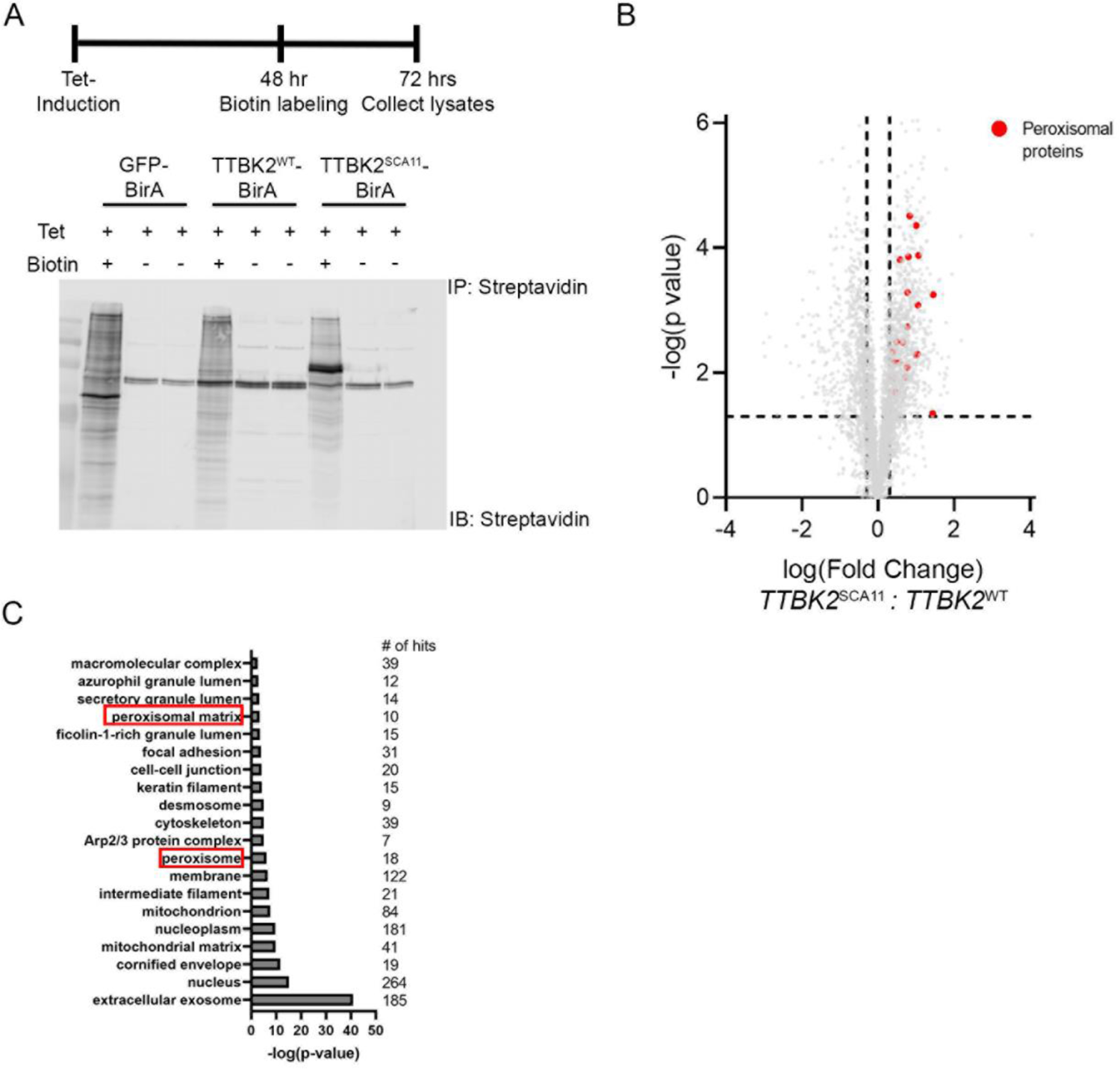
Comparative proximity proteomics identifies peroxisomal proteins as preferentially proximate to TTBK2^fam1^. (A) Schematic of BioID experimental design and test of biotin labeling. Flp-In tREx 293T cells were given Tet to induce expression of TTBK2^WT^- and TTBK2^fam1^-BirA. Two days later, Biotin was added to the media. Lysates were collected, pulled down using Streptavidin-conjugated beads, and blotted for Streptavidin. (B) Volcano plot depicting the interactome of TTBK2^fam1^ compared to TTBK2^WT^. Vertical dashed lines mark ±log (2) fold change. Horizontal dashed line marks P-value = 0.05. Red dots denote peroxisomal proteins. (C) Bar graph depicts the negative log value of GO terms of cellular localization and biological activity of proteins enriched in TTBK2^fam1^ BioID relative to TTBK2^WT^ BioID. The number of hits in each GO term is listed on the right.

**Figure S2.**
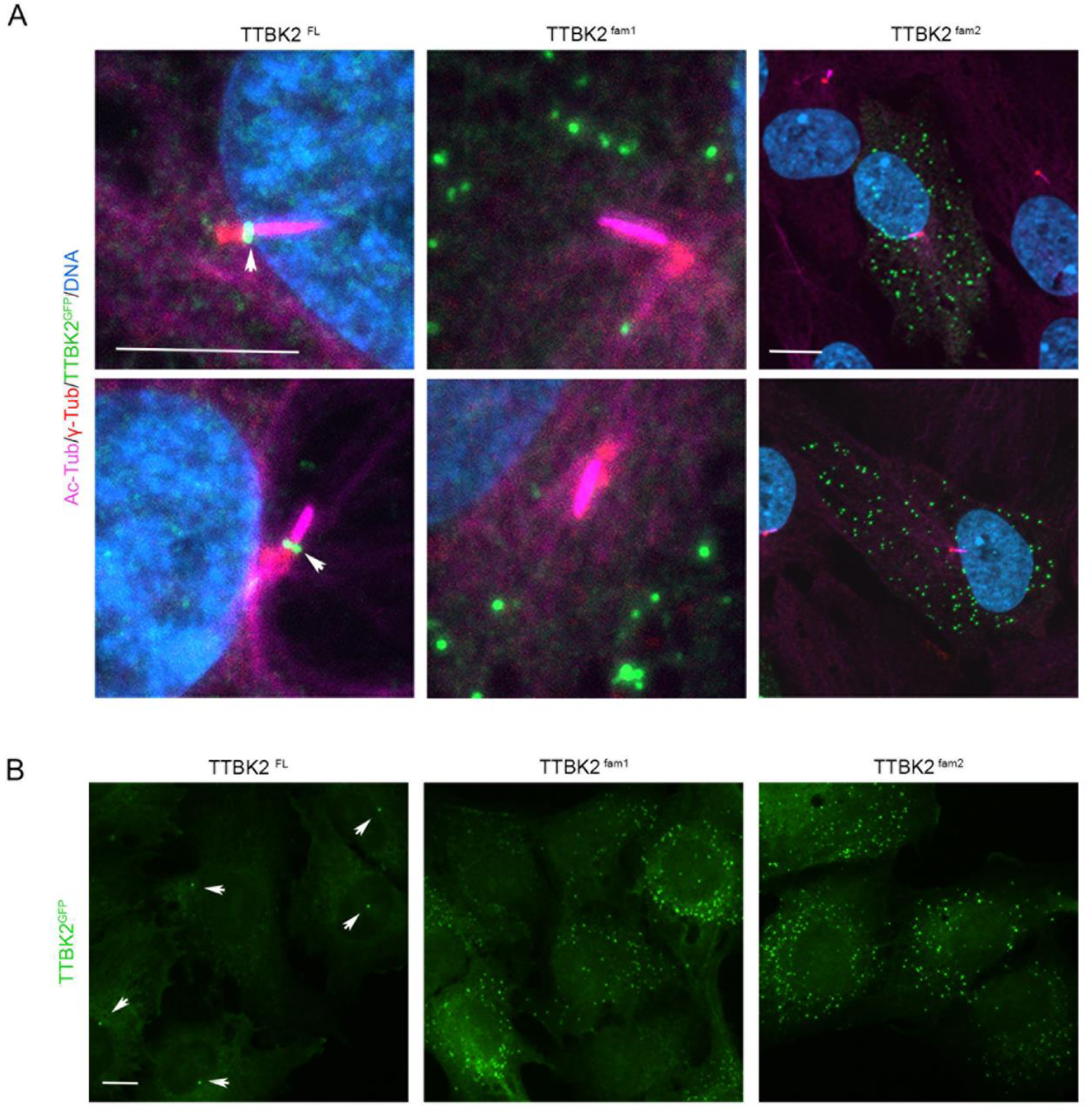
Frameshift mutations in *TTBK2* associated with SCA11 lead to loss of TTBK2 localization at the ciliary base. (A) Immunofluorescent images of RPE1 cells transiently expressing either full-length TTBK2 (TTBK2^FL^) or TTBK2 SCA11-causing proteins (TTBK2^fam1^ and TTBK2^fam2^) tagged with GFP-FLAG reporters. After cilia induction, cells were co-immunostained for Acetylated α-Tubulin (Ac-Tub, magenta), γ-Tubulin (γ-Tub, red) and GFP (green) to detect ciliary axoneme, basal body and subcellular localization of TTBK2 fusion proteins, respectively. DNA/nuclei (blue). Arrows depicts the characteristic localization of wild-type TTBK2 protein at the ciliary base. Bar, 5 μm and 10 μm (for TTBK2^fam2^). (B) Immunofluorescent images of TTBK2^FL^, TTBK2^fam1^ and TTBK2^fam2^ cells labeled with an antibody against GFP protein (green). Arrows indicate TTBK2 ciliary localization in TTBK2^FL^ cells, which is loss due to *TTBK2* mutations linked with SCA11 as observed in TTBK2^fam1^ and TTBK2^fam2^ cells. Bar 10 μm.

**Figure S3.**
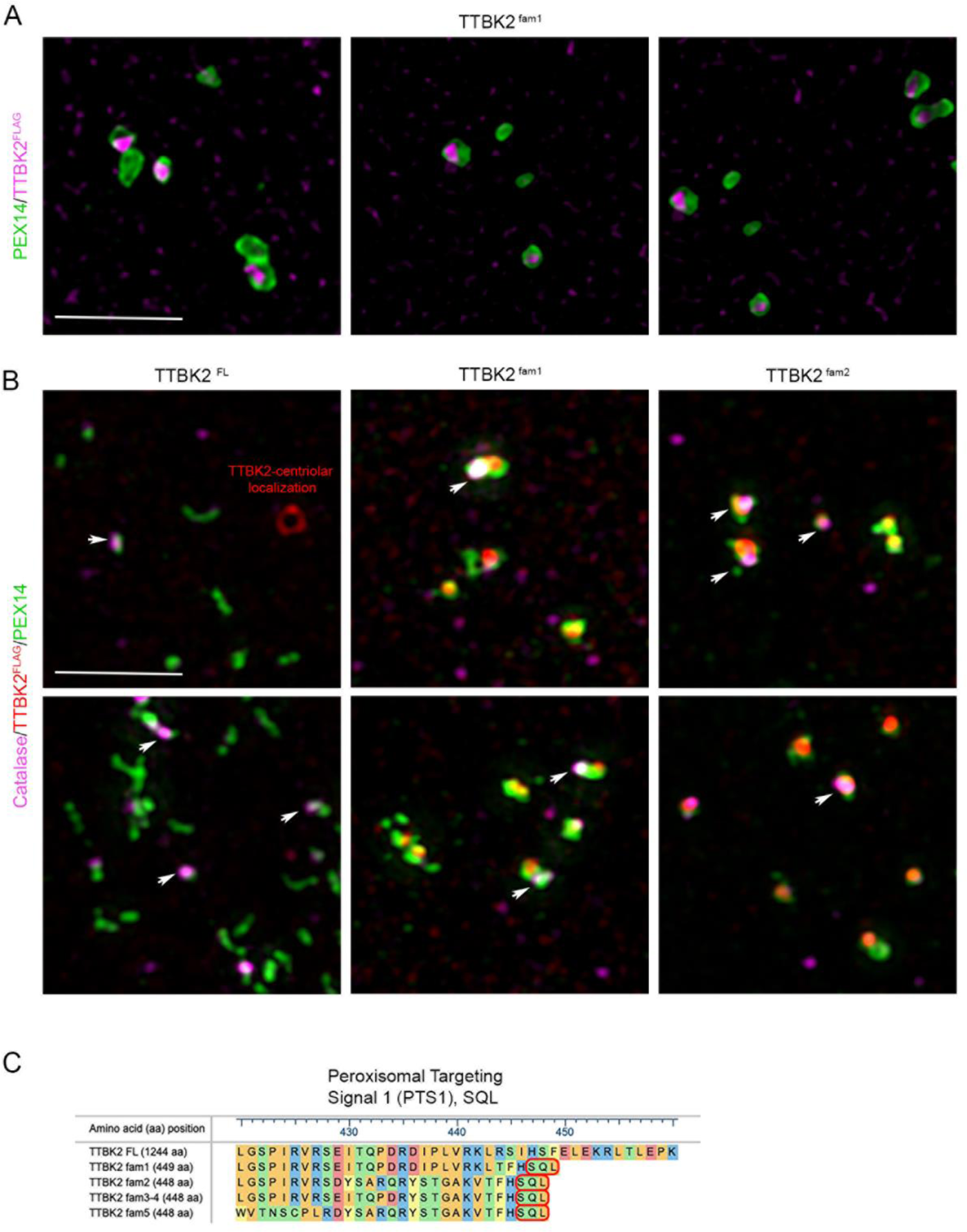
SCA11-causing truncated proteins are trafficked to the peroxisomal lumen. (A) Super-resolution fluorescent images of TTBK2^fam1^ cells co-immunostained for FLAG (magenta) and PEX14 (Proteintech, green) as peroxisome membrane marker. Bar, 2.5 μm. (B) Super-resolution fluorescent images of TTBK2^FL^ or TTBK2^fam1^ and TTBK2^fam2^ cells immunolabeled for catalase (R&D Systems, magenta), FLAG (red) and PEX14 (green). Catalase and PEX14 as matrix and membrane peroxisome markers. Arrows point out catalase localization to peroxisomes. Bar, 2.5 μm. (C) Scheme shows sequence alignment of amino acids (aa) of wild-type TTBK2 (420–460 aa) with the C-terminal domain of TTBK2 SCA11-causing proteins identified in 5 different family pedigrees worldwide. All *TTBK2* frameshift mutations are predicted to result in truncated protein products of ∼450 aa with a PTS1 consisting in the “Serine, glutamine and leucine” (or S, Q, L) motif.

**Figure S4.**
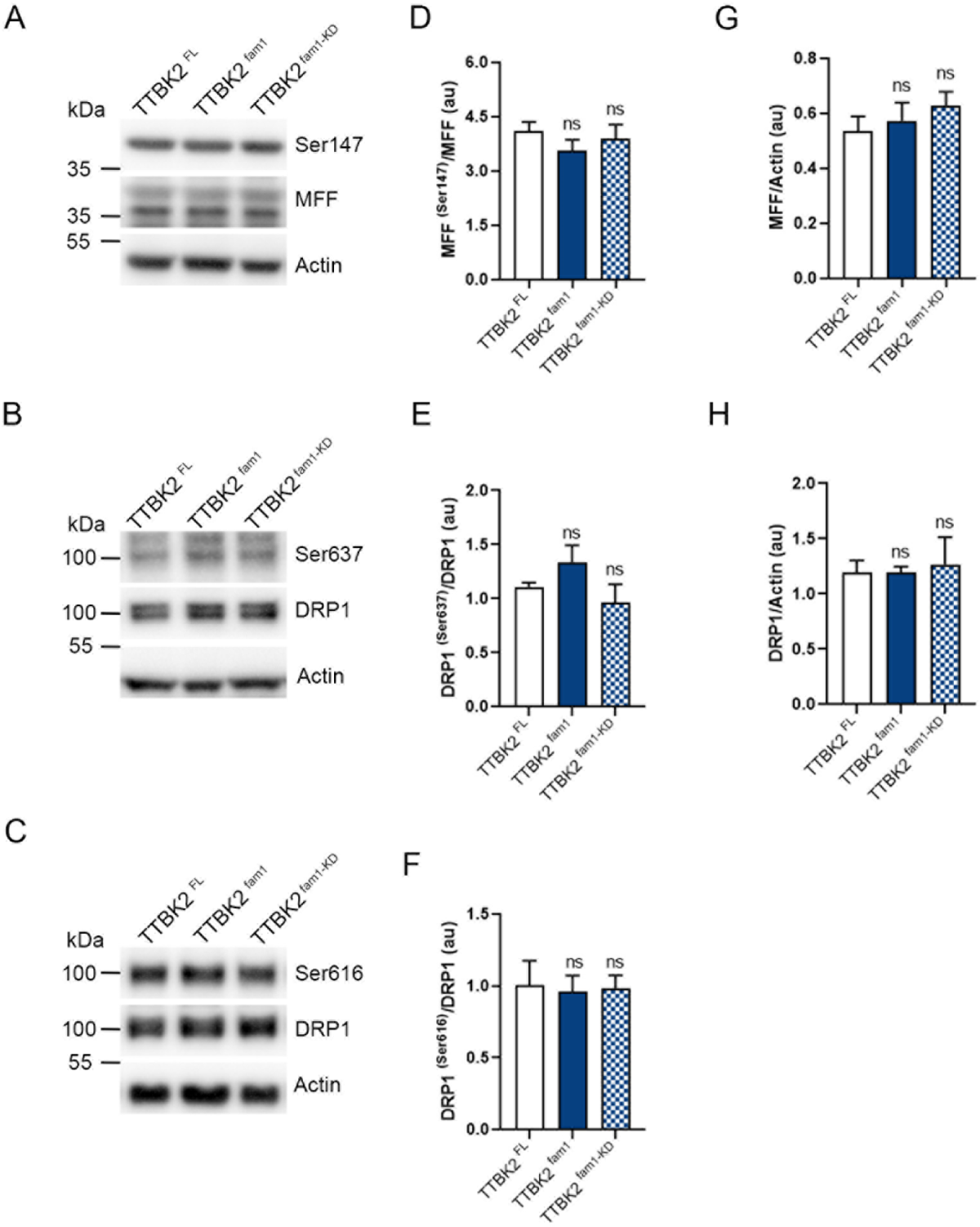
Serine phosphorylation analysis in MFF and DRP1 residues in cells expressing a SCA11-causing protein. (A-C) Immunoblotting analysis in TTBK2^FL^, TTBK2^fam1^ and TTBK2^fam1KD^ cell lysates with the indicated antibodies. Cells were lysate with NP40 buffer containing a phosphatase inhibitor cocktail (Cell Signaling). (A) Analysis of Ser147 for MFF (St John’s Lab) and (B, C) Ser637 and Ser616 for DRP1 (Cell Signaling). (D-F) Quantification of results by normalizing p-MFF/DRP1 to total MFF/DRP1 levels. (G, H) Quantification of relative MFF and DRP1 protein levels using actin for normalization. Bars represent mean ± s.e.m.: ns (not significant), one-way ANOVA with Dunnett’s multiple comparison test, n=4.

**Figure S5 1.**
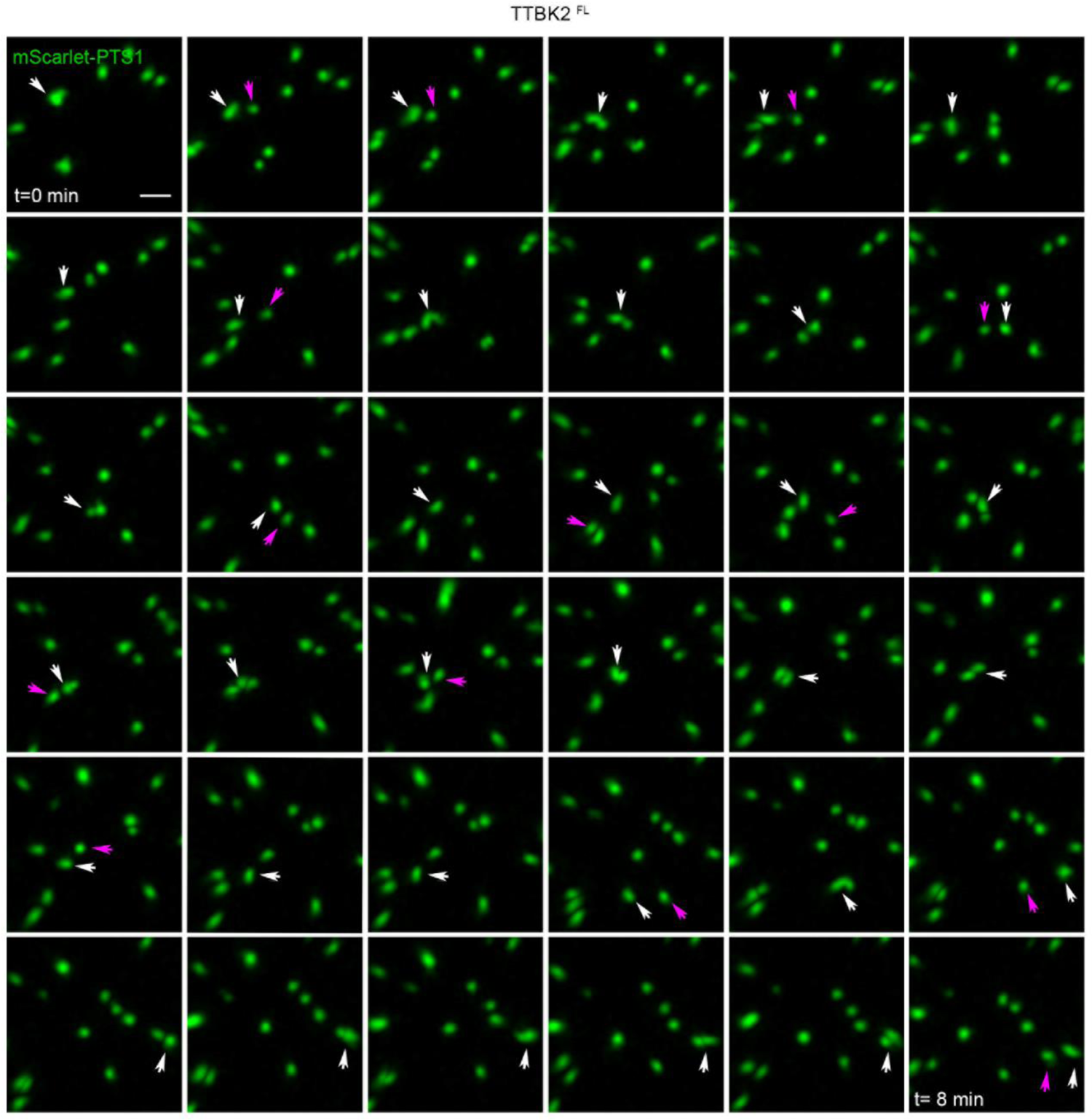
Time-lapse analysis of peroxisome dynamics in TTBK2^FL^ cells. Representative confocal images of peroxisome behavior over time within a region of a TTBK2^FL^ cell (**Video 1**). Peroxisomes were fluorescently labeled with mScarlet-PTS1 protein. White and magenta arrows depict the analyzed fluorescent particle and particle fusion events overtime, respectively. Bar, 10 μm.

**Figure S5 2.**
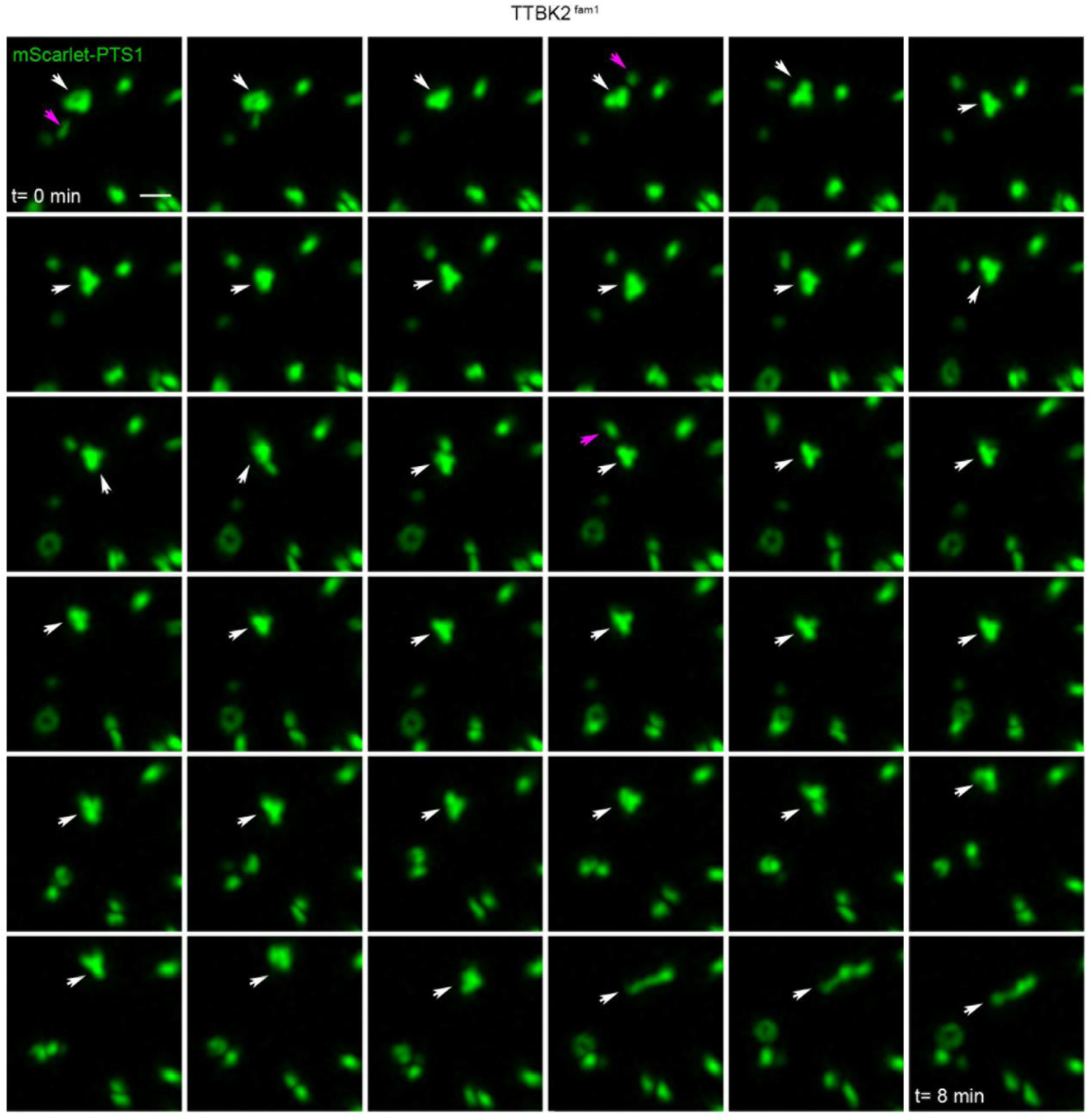
Time-lapse analysis of peroxisome dynamics in TTBK2^fam1^ cells. Representative confocal images of peroxisome behavior over time within a region of a TTBK2^fam1^ cell (**Video 2**). Peroxisomes were fluorescently labeled with mScarlet-PTS1 protein. White and magenta arrows depict the analyzed fluorescent particle and particle fusion events overtime, respectively. Bar, 10 μm.

**Figure S6.**
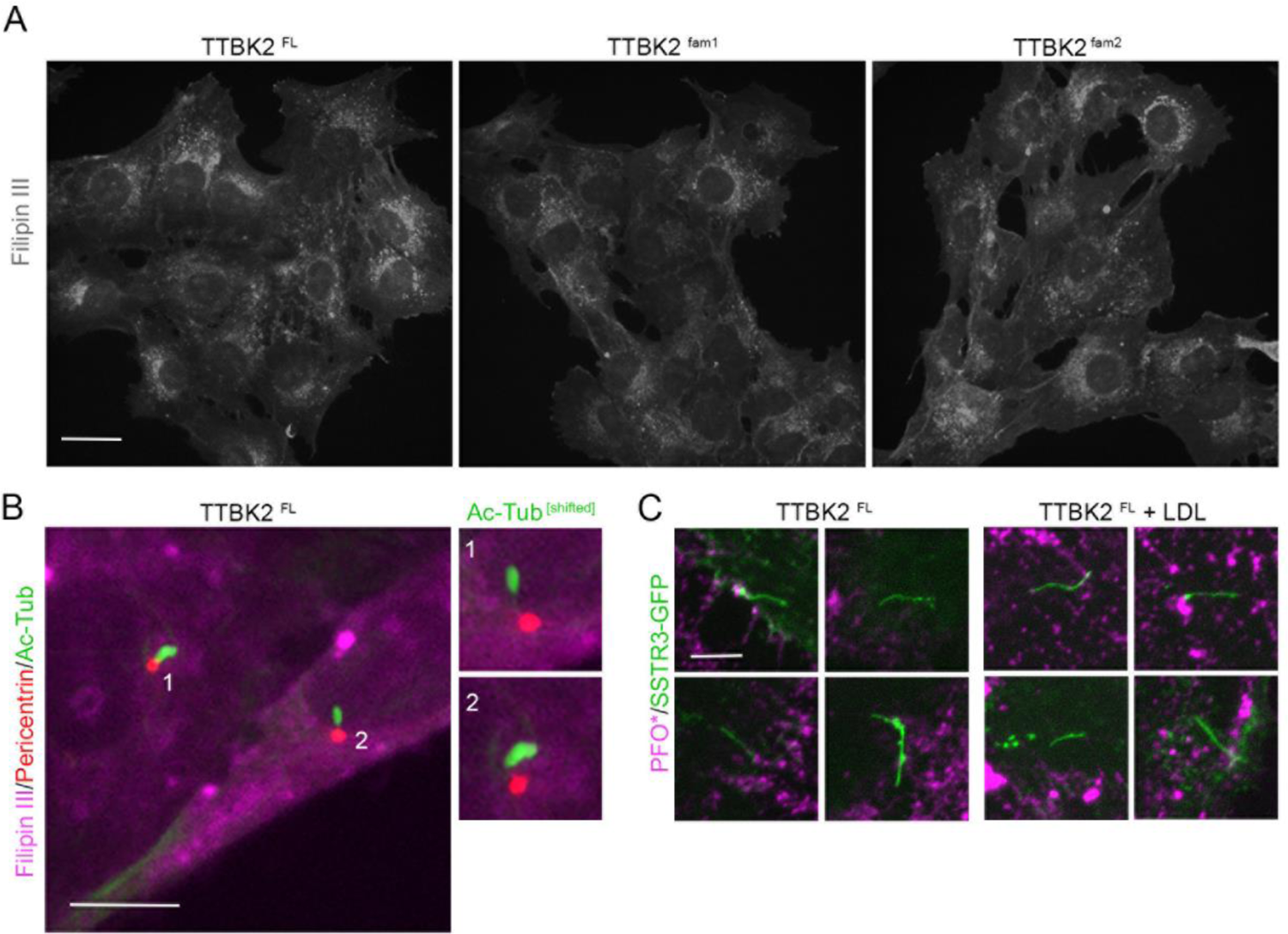
Detection of cholesterol in cells expressing SCA11-causing proteins. (A) Confocal images of cells expressing full-length TTBK2 (TTBK2^FL^) or SCA11-causing proteins TTBK2^fam1^ TTBK2^fam2^ stained with 50μg/ml of Filipin III fluorescent probe (R&D Systems). Unnoticeable differences in intracellular cholesterol content are observed among cell lines. Bar, 30 μm. (B) Fluorescent images of TTBK2^FL^ ciliated cells stained with 50 μg/ml Filipin III (magenta) and immunolabeled for pericentrin (red) and acetylated α-tubulin (Ac-Tub, green) as centrosome and cilia markers, respectively. Insets on the right show no evident presence of ciliary cholesterol. Bar, 10 μm. (C) Fluorescent images of TTBK2^FL^ ciliated cells transiently expressing ciliary somatostatin 3 receptor tagged with GFP (SSTR3-GFP) and fluorescently labeled with PFO* in the presence of 40 μM of myriocin (Sigma Aldrich) or (40 μM myriocin + LDL cholesterol). PFO* (or <u>P</u>erfringolysin <u>O</u>) binds accessible cholesterol and myriocin inhibits sphingomyelin synthesis, increasing accessible cholesterol (Kinnebrew et al., 2021). Undetectable cholesterol is observed at the ciliary membrane. Of note, detergent free conditions were used when testing fluorescent cholesterol-binding probes. Bar, 5 μm.

**Figure S7.**
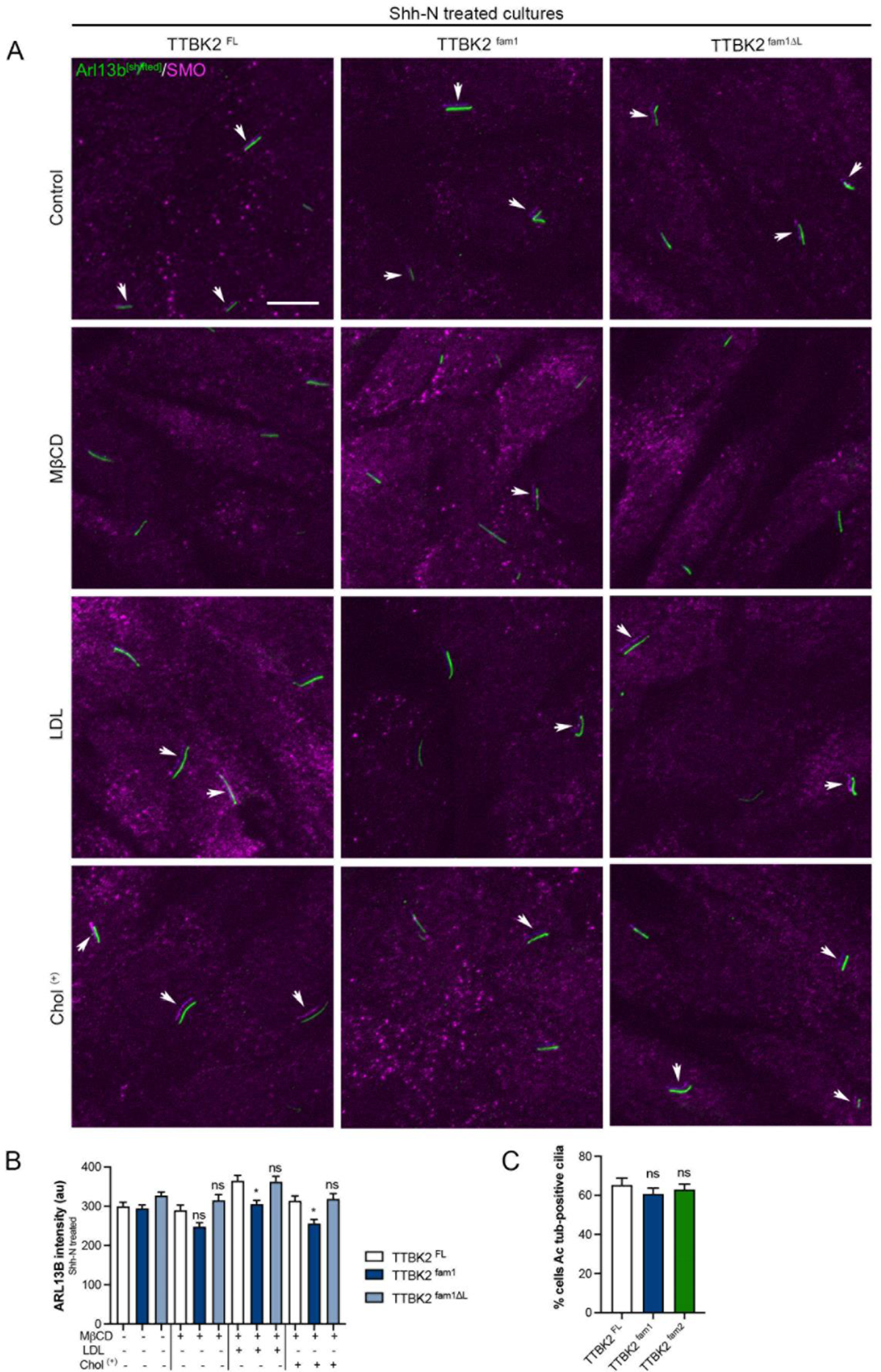
SCA11-causing proteins reduce trafficking of Alr13b to the ciliary membrane. (A) Immunofluorescent images of TTBK2^FL^, TTBK2^fam1^, TTBK2^fam1ΔL^ ciliated cells treated with 50 nM of SHH-N for 24 h (Control) and immunolabeled for SMO (magenta) and Arl13b (green) ciliary markers. Parallel cultures were depleted of cholesterol using Methyl-β-cyclodextrin (MβCD) and either supplemented with low*-density lipoprotein* cholesterol (LDL) or water-soluble cholesterol (Chol^+^) in the presence of 50 nM of SHH-N and 40 μM of cholesterol biosynthesis inhibitor pravastatin for 24h. Arrows point out SMO localization along the ciliary membrane upon SHH-N stimulation. Bar, 10 μm. (B) Quantification of Arl13b ciliary intensity in TTBK2^FL^, TTBK2^fam1^, TTBK2^fam1ΔL^ cell cultures with the indicated treatments. (C) Proportion of ciliated cells expressed as percentage TTBK2^FL^, TTBK2^fam1^, TTBK2^fam1ΔL^ using as a ciliary marker Acetylated α-Tubulin (Ac-Tub). Bars represent mean ± s.e.m.: ns (not significant), *P<0.05: Pearson’s chi-square (λ^2^) tests, n=2; 250 cells.

**Figure S8.**
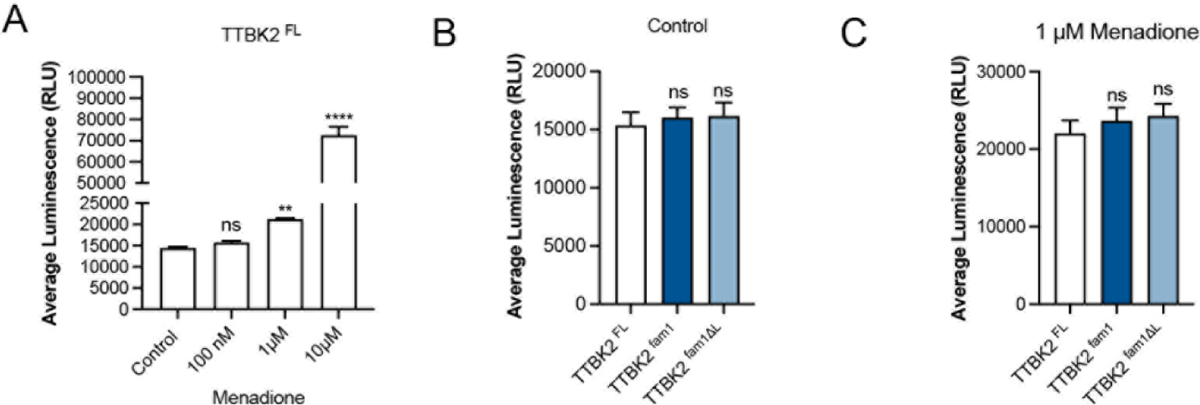
Determination of hydrogen peroxide (H_2_O_2_) levels in TTBK2^fam1^ cell cultures. (A) Relative luminescence units (RLU) of TTBK2^FL^ cells treated with increasing concentrations of ROS-induced compound menadione for 2 h. Levels of H_2_O_2_ were measured using ROS-Glo™ H_2_O_2_ Assay (Promega) according to manufacturer’s instructions. Relative luminescence units (RLU) were recorded using a plate reader. (B, C) RLU of TTBK2^FL^, TTBK2^fam1^, TTBK2^fam1ΔL^ control cell cultures or cultures treated with 1 μM of menadione (Sigma) for 2 h, respectively. Bars represent mean ± s.e.m.: ns (not significant), **P<0.01, ***P<0.001: Kruskal-Wallis test with Dunn’s multiple comparisons test, n=2; quadruplicates per experiment.

**Figure S9.**
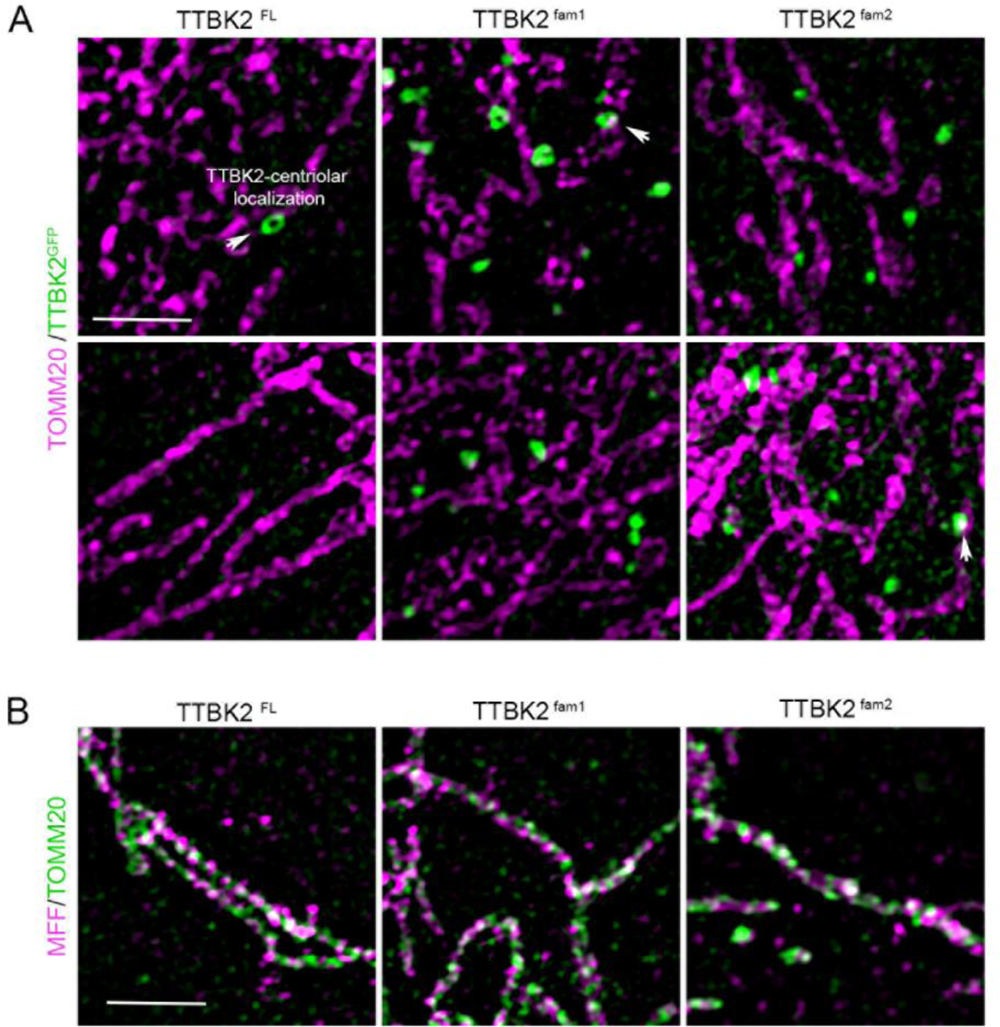
SCA11-causing proteins are not trafficked to mitochondria. (A) Super-resolution fluorescent images of TTBK2^FL^, TTBK2^fam1^ and TTBK2fam2 cells immunolabeled for translocase of outer mitochondrial membrane 20 (TOMM20, magenta) (Novus) and GFP (green). GFP detects localization of GFP-FLAG-fusion proteins. Images show wild-type TTBK2 centriolar localization in TTBK2FL cells (arrowhead) and peroxisomal localization of SCA11-associated proteins in TTBK2^fam1^ and TTBK2^fam2^ cells (speckle staining pattern). The slight overlapping of GFP fluorescent signals with TOMM20 in TTBK2^SCA11^ cells may be indicative of the recognized membrane contact between peroxisomes and mitochondria (arrows). Bar, 2.5 µm. (B) Representative fluorescent images of MFF (green) and TOMM20 (green) in TTBK2FL, TTBK2fam1 and TTBK2fam2 cells. Not evident accumulation of MFF is noticed at mitochondria membranes as observed for peroxisomes (**Fig 4A**). Bar, 2.5 µm.

